# Causal contribution of motor-parietal network to conscious kinesthetic perception

**DOI:** 10.64898/2026.10.07.757148

**Authors:** Ozge Ozlem Saracbasi, Tomoyo Morita, Hideki Nakano, Yandi Tang, Yuki Sato, Jihoon Park, Eiichi Naito

## Abstract

Bodily awareness, including the conscious experience of limb movements (kinesthesia), is a fundamental form of human consciousness. Tendon vibration–induced kinesthetic illusions provide a tractable model for probing its neural basis. Although the primary motor cortex (M1), right inferior parietal lobule (IPL), and right inferior frontal gyrus (IFG) are coactivated during illusory movement, whether and how the coordinated engagement of modality-specific motor and higher-order cortical regions causally contributes to conscious bodily experience remains unclear. Here, we combined functional magnetic resonance imaging (fMRI) and transcranial direct current stimulation (tDCS) to investigate the neural mechanisms and causal contribution of these regions to conscious kinesthetic perception. fMRI revealed that M1–IPL network supports kinesthetic illusions emerging through bottom-up sensory processing, whereas the IFG was selectively recruited when previously unnoticed kinesthetic signals became consciously experienced following knowledge, consistent with a role in top-down monitoring. Causal analysis further revealed a consistent bottom-up directional relationship from M1 to the IPL. To test its causal contribution, we applied concurrent facilitative dual-site tDCS to M1 and the IPL. M1–IPL tDCS selectively enhanced the subjective strength of the illusion compared with M1–IFG and sham stimulation. This effect was replicated in an independent cohort and exceeded the effects of stimulating M1 or the IPL alone and the IPL–IFG, without detectable muscle activation. These findings provide causal evidence that coordinated motor–parietal engagement contributes to conscious kinesthetic experience and provide new insights into how conscious sensory perception can be enhanced through interactions between modality-specific and higher-order cortical systems.

**Significance statement:** Bodily awareness is fundamental to human consciousness, yet how cortical networks transform bodily signals into conscious experience remains unclear. Using tendon vibration-induced kinesthetic illusions, we combined functional magnetic resonance imaging (fMRI) and dual-site transcranial direct current stimulation (tDCS) to distinguish bottom-up from top-down processes and test their causal contributions. fMRI implicated the primary motor cortex (M1)–right inferior parietal lobule (IPL) network in bottom-up kinesthetic processing, whereas the right inferior frontal gyrus was selectively recruited during top-down monitoring. Concurrent M1–IPL tDCS robustly and reproducibly enhanced conscious kinesthetic perception without detectable muscle activation, exceeding other stimulation configurations. These findings provide causal evidence that coordinated motor–parietal engagement contributes to conscious bodily experience and advance mechanistic understanding of bodily awareness.

## Introduction

Understanding how neural activity gives rise to conscious experience remains a fundamental challenge in neuroscience. Among various forms of consciousness, bodily awareness—the perception of one’s own body and its movements—is fundamental to human consciousness. A key component of bodily awareness is kinesthesia, the conscious perception of limb movement. While kinesthetic signals from muscle spindles elicit reflexive and unconscious motor responses via the spinal cord and the primary motor cortex (M1), these same signals can elicit vivid kinesthetic illusion—a conscious sensation of movement without actual motion—when tendon vibration is applied (1, 2, 3). For example, tendon vibration of the tibialis anterior (TA) muscle can induce an illusory sensation of plantarflexion of the relaxed foot (4). In this way, kinesthetic illusion provides a unique model for investigating the neural mechanisms of bodily awareness because conscious kinesthetic perception can be generated without actual movement.

Neuroimaging studies have consistently identified a cortical network associated with kinesthetic illusions, including the primary motor cortex (M1), the right inferior parietal lobule (IPL), the right inferior frontal gyrus (IFG), and the bilateral insular cortices (4–11). Among these regions, lesion in M1 impairs the illusion (12), and damage to white matter fibers connecting the right IPL and IFG disrupts proprioception (13). In addition, activity in the right IPL and IFG increases during kinesthetic illusion and correlates with illusion strength (9, 14). These findings suggest that interactions among the M1, the right IPL, and the right IFG are critical for generating conscious kinesthetic perception.

However, the functional organization of this network remains unresolved. In particular, it is unclear whether the right IPL and IFG make distinct contributions to kinesthetic awareness, though these regions are strongly interconnected (14–16). In general, the IPL has been proposed to support higher-order integration of bodily sensory information (8), whereas the right IFG, including the ventral premotor cortex, has been implicated in monitoring sensorimotor states (17, 18). Such functional differences suggest that these regions may play respective roles in conscious kinesthetic perception.

The previous studies of kinesthetic illusions (see above) have typically required participants to attend to or expect the illusory sensation during tendon vibration. Such top-down instruction, however, makes it difficult to distinguish neural processes that directly generate conscious kinesthetic perception from those involved in monitoring or expectation. In the case of a visual illusion, without instructions, some participants spontaneously experience the illusion, whereas others do not; however, after receiving explicit instruction, participants notice it (19). We have empirically known that similar phenomenon also occurs particularly with kinesthetic illusion of foot movement (4). Therefore, manipulating prior knowledge of vibration-induced illusion provides an approach to dissociate bottom-up sensory processes from top-down influences on bodily awareness.

In the present functional magnetic resonance imaging (fMRI) study, we investigated the functional organization of the network of the M1, the right IPL, and the right IFG and their respective roles during kinesthetic illusion. Tendon vibration was applied to the TA muscle of the relaxed right foot to induce illusory plantarflexion. In order to experience kinesthetic illusion during tendon vibration, participants must be aware of movement sensations amidst the vibratory sensations. Participants first completed two sessions without being informed that tendon vibration could produce illusory movement (Sessions 1 and 2), followed by two sessions after receiving this information (Sessions 3 and 4). We hypothesized that illusions experienced without prior knowledge would be primarily elicited by bottom-up kinesthetic processing, whereas illusions experienced after receiving the knowledge would additionally involve top-down mechanisms. We first show that illusion with knowledge recruits the previously identified kinesthetic network, but the right IPL and the right IFG make distinct contributions to the illusion. We further show functional coupling and causal relationship among the M1, the right IPL, and the right IFG. By doing so, we provide evidence for a distributed cortical mechanism underlying conscious kinesthetic perception.

Conscious perception across sensory modalities (visual, auditory, and tactile) has been proposed to emerge through interactions between modality-specific sensory regions and distributed frontoparietal networks, as described by the Global Neuronal Workspace (GNW) hypothesis (20–24). However, direct causal evidence that such interactions generate conscious perception remains limited. Kinesthesia provides an opportunity to test this principle in the domain of bodily awareness, as our fMRI results identified coactivation of the modality-specific kinesthetic-motor cortex (M1) and higher-order right frontoparietal regions (IPL and IFG) during kinesthetic illusions. We therefore tested whether facilitating these regions with dual-site transcranial direct current stimulation (tDCS) could causally modulate kinesthetic awareness. M1–IPL tDCS enhanced kinesthetic illusion strength more than M1–IFG tDCS, and this effect was replicated without increasing muscle activity and exceeded the effects of single-site M1 or IPL tDCS and dual-site IPL–IFG tDCS. These findings provide causal evidence that coordinated interactions between the M1 and the right IPL contribute to conscious kinesthetic awareness, extending the GNW framework to bodily awareness.

## Results

### fMRI experiment

#### Aware and unaware of illusion

We recruited a total of 62 healthy younger adults in the fMRI experiment, who did not have prior knowledge about vibration-induced illusory movement before the experiment. We conducted four sessions in the experiment. In the fMRI scanner, we vibrated the tendon of tibialis anterior (TA) muscle of their relaxed right foot (Figure 1A), without giving them prior knowledge of vibration-induced illusory movement (sessions 1 and 2 referred to as s12), and after giving them the knowledge (sessions 3 and 4 referred to as s34). In s12, 22 (20 women) of 62 participants reported that they experienced sensation of plantarflexion of their stationary right foot even without the knowledge (Figure 1B). In s34, in addition to these 22 participants, 30 participants (20 men) who had not noticed the illusion in s12 became aware of and experienced the illusion after receiving the knowledge. Three participants (two men) did not explicitly report the illusion in s12, though they reported that they might feel it retrospectively during s12. However, we determined that the reliability of their responses was low and therefore excluded these participants from the subsequent grouping. Seven participants (four men) reported no illusions throughout the entire experiment, and were excluded from the analysis (Figure 1B).

**Figure 1.**
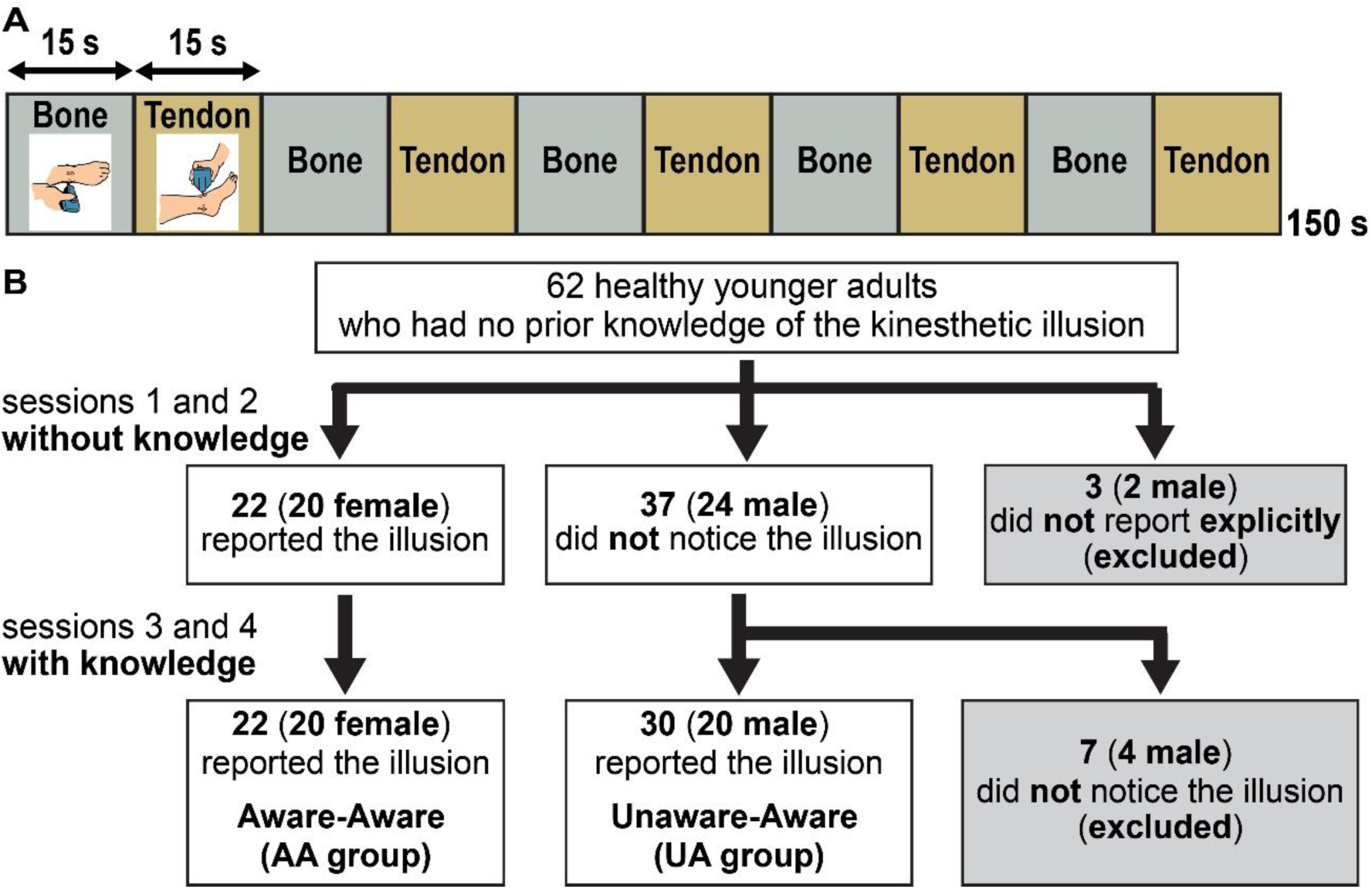
Illustration of fMRI experimental procedure. (A) Task design. Each participant completed four sessions, each including five 15-sec tendon-vibration and five 15-sec bone-vibration epochs, 150-sec in total. (B) Flowchart of participant grouping.

The 22 participants who explicitly reported the illusion both with and without knowledge were assigned to aware-aware group (AA group; Figure 1B), whereas the 30 participants who did not experience the illusion without knowledge but did experience it with knowledge was classified unaware-aware group (UA group; Figure 1B). The majority of participants in the AA group were women (20 of 22), whereas the majority of participants in the UA group were men (20 of 30). The sex distribution differed significantly between the two groups (χ^2^(1) = 17.2, p < 0.001, see SI Appendix). Figure S1 shows maximum illusory angle reported during s12 and during s34 in the UA and AA groups (SI Appendix).

#### Brain activation, functional connectivity, and causal relationship

We measured brain activity when we vibrated the tendon of TA muscle of their relaxed right foot during a task (Tendon) epoch, and while we vibrated the skin surface over the lateral malleolus of fibula of the right foot during resting period (Figure 1A). Tendon vibration-induced illusion-related activity was tentatively determined as an activity increase during the Tendon epoch from that during the resting (Bone) period. We first conducted a full factorial design analysis: a within-subject factor (session [2]: s12, s34) and a between-subject factor (group [2]: AA, UA).

When both AA and UA groups reported the illusion during the last two sessions (s34), illusion-related activity was observed in the foot section of the left M1, the bilateral SMA, the right IPL, the right IFG, and the bilateral insular cortices (Figure 2A and SI Appendix Table S1) within the regions-of-interest defined by a study in which we depicted brain regions showing the right-foot illusion-related activity (Tendon > Bone) during illusion with knowledge in an independent group of 57 healthy participants (SI Appendix and Table S2).

**Figure 2.**
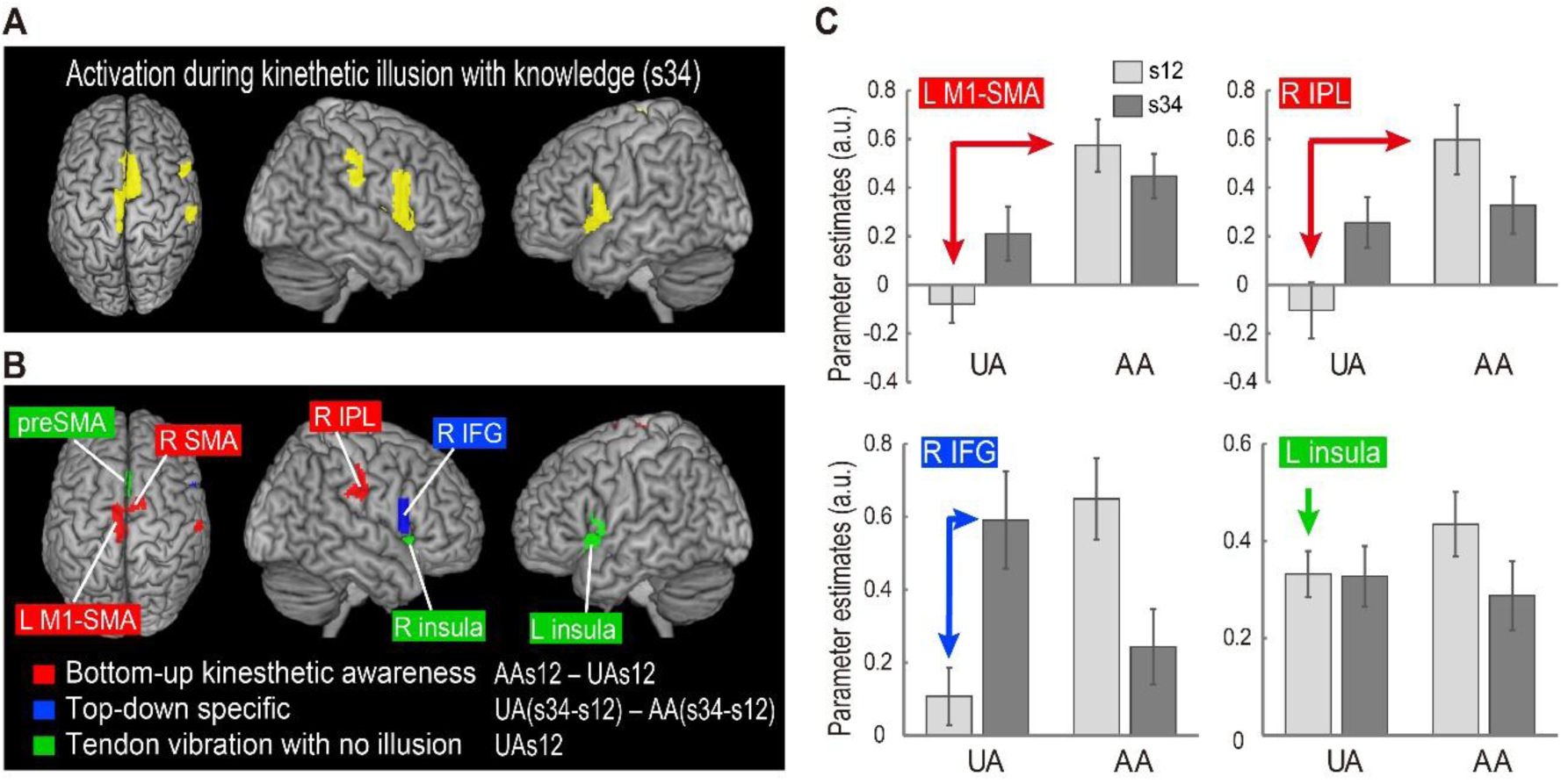
Results of fMRI contrast analysis. (A) Brain regions showing significant activation during kinesthetic illusion with knowledge. (B) Red: Brain regions active when AA group experienced the illusion without knowledge in a bottom-up manner as compared to when UA group did not. Blue: Right IFG region active when UA group noticed and experienced the illusion while attending it, compared with when they did not experience it without knowledge, by controlling for the session order effect. Green: Brain regions active during tendon vibration even when UA group did not experience the illusion. Details of the anatomical locations of brain activations are provided in SI Appendix Tables S1, S3 and S4. (C) Averaged brain activity (parameter estimate) across participants in UA and AA groups. The activity was extracted from the significant clusters identified in panel B for each participant, and the average was calculated. Error bars indicate standard errors of the mean across participants.

Next, by comparing brain activity during s12 between the AA and UA groups, we identified brain regions showing activity increase when the AA group experienced the illusion without knowledge compared with when the UA group did not. Significant clusters of voxels were identified in the foot section of left M1 (area 4a), the bilateral SMA, and the right IPL (area PFt; Figure 2B red sections and SI Appendix Table S3), suggesting that these are important regions when participants experience the illusion in a bottom-up manner. When we looked at the activity in each region, we confirmed activity increase in the AA group, whereas no substantial increase in the UA group during s12 (Figure 2C top two panels).

We further examined brain regions in which activity increased selectively when the UA group noticed and experienced the illusion while attending it, compared with when they did not, by controlling for the session order effect, i.e. interaction term [s34 vs. s12 in UA group vs. s34 vs. s12 in AA group]. We found significant activity increase (p = 0.05 FWE-corrected) only in the right IFG (area 44; Figure 2B blue sections and SI Appendix Table S4), suggesting this region increases its activity selectively when participants notice and experience the illusion by engaging in top-down monitoring of kinesthetic signals to expect and predict illusory movement during tendon vibration.

Increased activity (Tendon > Bone) was also observed in the bilateral anterior insular cortices, extending into the bilateral putamen (Figure 2B green sections). However, unlike the M1, SMA, IPL, and IFG, the bilateral insular cortices were also activated during tendon vibration even when the UA group did not experience the illusion (s12 in UA group; Figure 2C right bottom). Such activation was also observed in the preSMA (Figure 2B green section). This suggests that the insular cortices responded to the change in stimulation from bone to tendon rather than being specifically involved in the illusion itself. Therefore, the M1, SMA, right IPL, and right IFG are likely to play key roles in experiencing the illusion.

Next, we conducted functional connectivity analyses to examine the functional coupling among the M1, SMA, right IPL and right IFG during the illusion with knowledge, using each region as a seed. We examined changes in functional connectivity when the UA group noticed and experienced the illusion with knowledge compared with when they did not, while controlling for the session order effect, using the interaction contrast [(s34 vs. s12 in UA group) × (s34 vs. s12 in AA group)]. Only when the M1 served as the seed did the right IPL (areas PF and PFcm) and the right IFG (area 44) showed increased functional connectivity with it (Figure 3A; SI Appendix Table S5). No regions showed increased functional coupling when the left or right SMA, the right IPL, or the right IFG was used as the seed. These findings indicate that the network involving the M1, the right IPL and the right IFG plays a particularly important role in kinesthetic illusion with knowledge.

**Figure 3.**
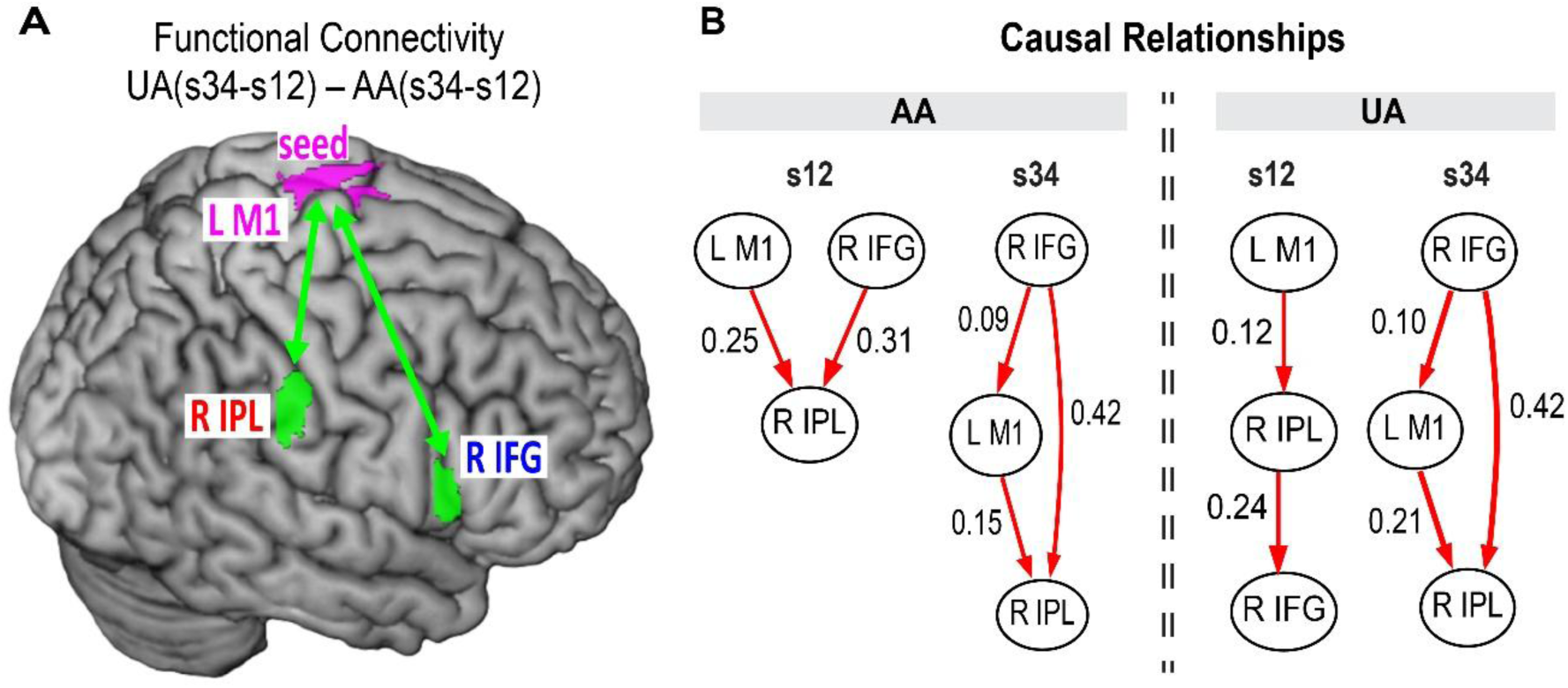
Results of functional connectivity and causality analyses. (A) The right IPL and IFG regions (green sections) showing enhanced functional coupling with the left M1 (seed, magenta section). (B) Significant causal relationships among ROIs (left M1, right IPL, right IFG). Arrows indicate causal effects from one ROI to another, and numerical values represent causal inference values.

The functional connectivity analysis does not reveal causal relationships (ordering) across brain regions. Therefore, we conducted a Linear Non-Gaussian Acyclic Model (LiNGAM) to explore the structural causal relationships among the three regions (the left M1 foot section, the right IPL and the right IFG) in the AA and UA groups. We conducted DirectLiNGAM for the first or last two sessions in the AA and UA groups separately (Figure 3B). In the last two sessions (s34) where both groups experienced the illusion with knowledge, the causal orders among the three regions were the same between two groups. The right IFG was the top of the causal hierarchy giving positive direct inference both to the M1 and to the right IPL, suggesting top-down function of the right IFG may facilitate neural processing of the M1-IPL bottom-up network. The M1 also gave a positive direct inference to the IPL. These causal structures were distinct when the AA group experienced the illusion without knowledge (s12); both the right IFG and the M1 gave positive direct inference to the right IPL independently without direct inference from the right IFG to the M1. Notably, direct positive inference from the M1 to the IPL was consistently observed no matter when the AA and UA groups experienced the illusion with or without knowledge and even when the UA group experienced no illusion without knowledge, suggesting that this is a fundamental causal relationship between the M1 and the right IPL.

#### tDCS experiments

Prompted by the enhanced functional coupling of the right IPL and the right IFG with the left M1 and the causal relationships among these regions during kinesthetic illusion with knowledge (Figure 3), we conducted a double-blind, parallel, randomized, sham-controlled tDCS study. We first examined whether concurrent facilitation of the left M1 and the right IPL (M1–IPL) and of the left M1 and the right IFG (M1–IFG) may causally augment conscious kinesthetic perception by applying dual-site facilitative tDCS to the foot section of the left M1 and the right IPL or IFG, respectively (Figure 4). Given that susceptibility to illusions appears to exhibit sex differences in the fMRI experiment (Figure 1), only male participants were recruited for the following tDCS experiments. We recruited new male participants and collected 63 UA type participants. Twenty-one of them were randomly assigned to M1-IPL, M1-IFG, or sham group, respectively. Before the tDCS experiment, we also measured brain activity during illusion with knowledge in all participants using fMRI (see SI Appendix), and confirmed that the illusion activated the highly similar regions as observed in the fMRI experiment (Figure 5A).

**Figure 4.**
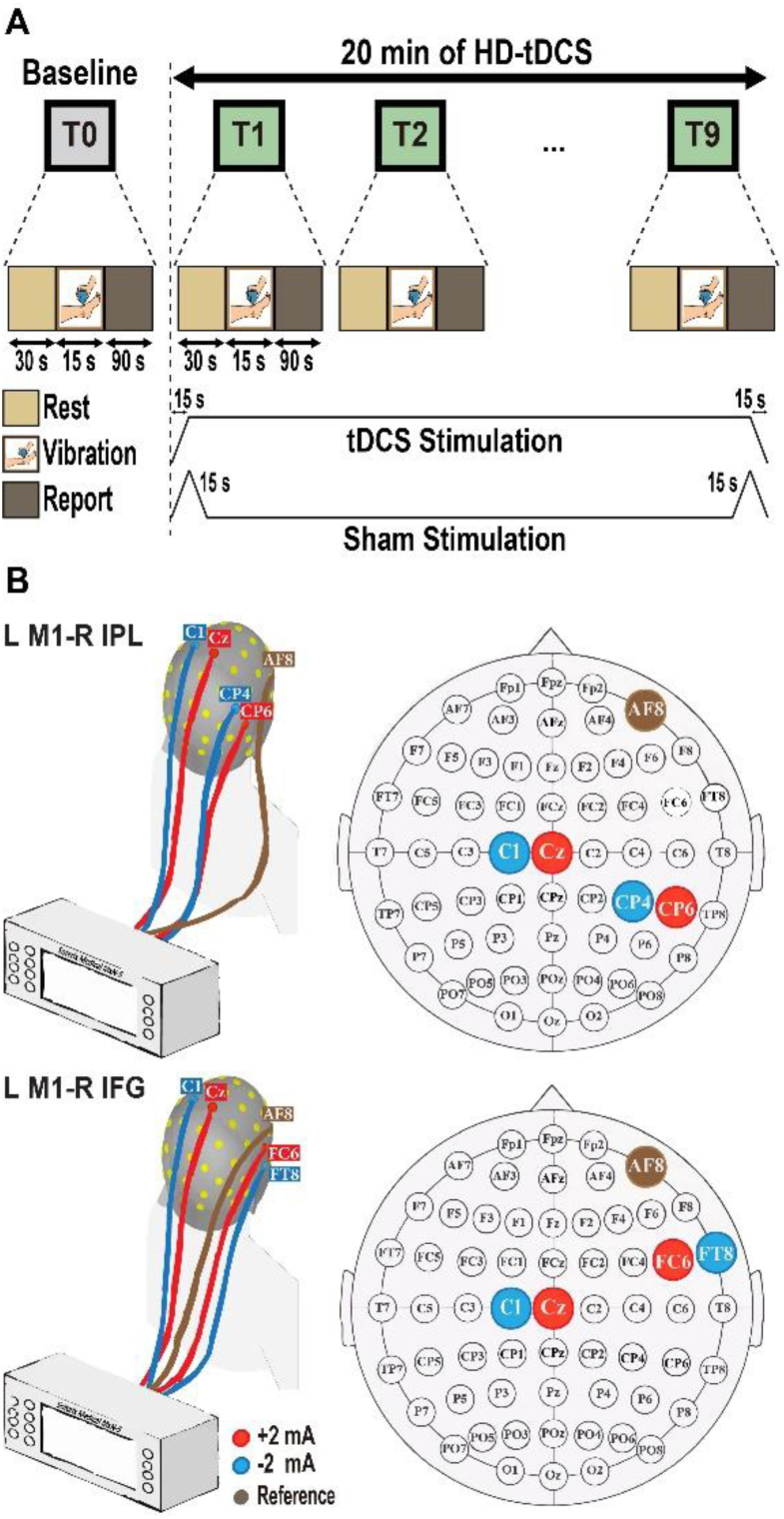
Schematic illustration of tDCS experimental design and electrode locations. (A) Experimental design and procedure of real tDCS and sham stimulation. (B) Left panels illustrate the experimental set-up for each of concurrent dual-site facilitative tDCS targeting to the left M1 and the right IPL (top) and to the left M1 and the right IFG (bottom). Right panels show the electrode locations based on the 10–20 international EEG system. Cz: anode and C1: cathode for left M1 (L M1), CP6: anode and CP4: cathode for right IPL (R IPL), FC6: anode and FT8: cathode for right IFG (R IFG). Reference electrode was placed at AF8.

**Figure 5.**
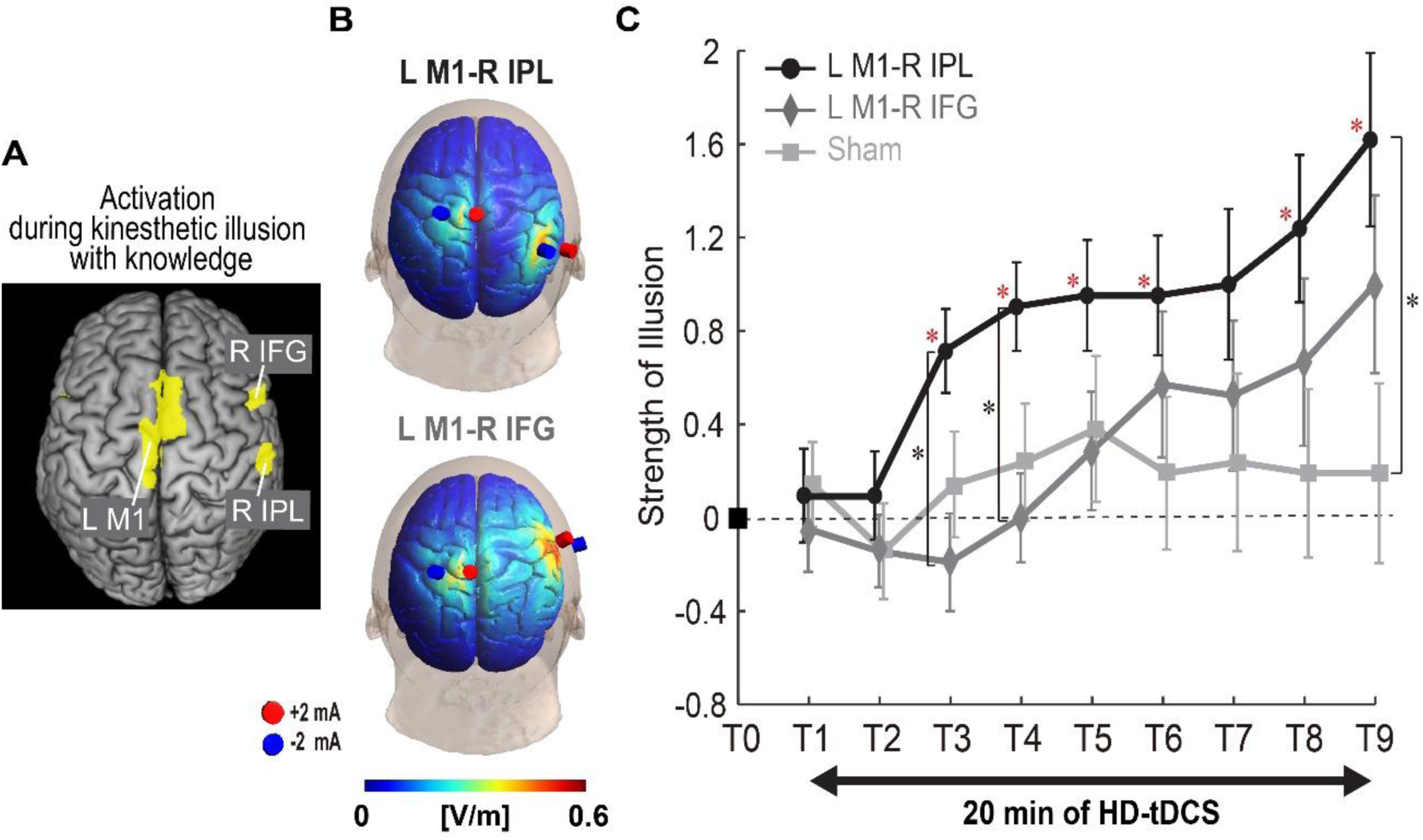
Illusion-related activity, simulated electric field, and tDCS results. (A) Illusion-related activity (group effect) in the left M1, the right IPL and the right IFG during illusion with knowledge. (B) Electric field simulation (group effect) using SimNIBS version 4.5.0 in M1–IPL and M1–IFG groups; red: anode, blue: cathode. Distribution of electric field (EF) is simulated based on average electrode locations across participants and is displayed on the MNI standard brain. The EF strength is visualized using a color scale ranging from blue to red. (C) Change in strength of illusion during 20-min tDCS. Red asterisks indicate significant difference when each trial (T1 to T9) was compared with baseline trial (T0) after Bonferroni correction. Black ones indicate significant between-group difference in each trial (corrected p < 0.05). MNI: Montreal Neurological Institute.

In the tDCS experiment, a total of ten 15-s tendon vibration trials were conducted (Figure 4A). Starting a baseline trial with no tDCS (T0), tDCS was provided for 20 min during which nine subsequent vibration trials were performed. In sham group, tDCS was provided only for the first and last 30-s (15-s ramp-up and 15-s ramp-down). In each trial, the participants were asked to attend to illusory plantarflexion of the right relaxed foot, and to report whether illusion experienced in a trial becomes stronger (+1), almost the same (0) or weaker (−1) than that experienced in one previous trial, and we simply added the scores across participants.

The participants could not identify the stimulation they received above chance level (50%), indicating that the blinding was effective (see SI Appendix). After the experiment, we confirmed that the target regions activated during the illusion (Figure 5A) could be effectively stimulated by simulating the distribution and strength of the electric field generated by the tDCS (Figure 5B, SI Appendix and Figures S5 and S6A, B). Prominent increase in the strength of the illusion was observed in the M1–IPL group (Figure 5C).

When we performed one-sample t-test, the illusion strength was significantly greater after the 3^rd^ trials (except 7^th^) than that in the baseline trial (T0) only in the M1–IPL group (p = 0.026, 0.004, 0.024, 0.045, 0.028, and 0.011 after Bonferroni correction, respectively). No such significant augmentation was observed in the M1–IFG and sham groups, suggesting prominent augmentation effect of illusion only in the M1–IPL group. Next, we examined difference across the M1–IPL, M1–IFG and sham groups. A 2-factor mixed-design ANOVA (group (3) x trials (9)) showed significant trend of interaction (F(16, 480) = 2.550, p = 0.087, partial η² = 0.078). Post hoc analysis revealed that the M1–IPL group reported significantly greater strength of illusion in the 3^rd^ and 4^th^ trials than the M1–IFG group (p = 0.011, 0.014), and that the illusion strength in the last (9^th^) trial (T9) in the M1–IPL group was significantly greater than that in the sham group (p = 0.035), which was not observed in the M1–IFG group (Figure 5C).

We also conducted a second double-blind, parallel, randomized tDCS study in an independent cohort of participants. We recruited a new sample of male participants and identified 75 UA-type participants. Fifteen participants were randomly assigned to each of five stimulation groups: M1–IPL, M1–alone, IPL–alone, IPL–IFG, and IFG–alone. The aims of this second tDCS study were to (I) examine the replicability of the augmentation effect of the M1–IPL tDCS, and (II) evaluate the effect of concurrent dual-site stimulation by comparing the M1–IPL tDCS with M1–alone and IPL–alone tDCS. We also investigated (III) whether concurrent dual-site stimulation of the two higher-order regions, the right IPL and the right IFG, would produce an augmentation effect comparable to that of the M1–IPL tDCS. Finally, based on the causality analysis (Figure 3B), which identified the IFG as the highest node in the causal hierarchy, exerting positive direct influences on both the M1 and the IPL, (IV) we compared the effects of the M1–IPL tDCS with those of IFG–alone tDCS.

Prominent increase in the strength of the illusion was replicated in the M1–IPL group (Figure 6A). In each group, we first examined augmentation of illusion in trials (T1 to T9) during tDCS as compared to the baseline trial (T0). One-sample t-test revealed that the illusion was significantly augmented after the 3^rd^ trials (except 5th) only in the M1–IPL group (p = 0.013, p < 0.001, p = 0.004, p < 0.001, p = 0.008 and p = 0.001 after Bonferroni correction, respectively; Figure 6), as observed in the first tDCS study (Figure 5C). None of the other groups (M1–alone, IPL–alone, IPL–IFG, and IFG–alone) showed significant augmentation. Next, we examined difference across the M1–IPL, M1–alone, and IPL–alone groups. A two-factor mixed-design ANOVA (group [3] x trials [9]) showed significant trend of interaction (F(16, 336) = 2.808, p = 0.072, partial η² = 0.118). The M1–IPL group reported significantly greater illusion strength in the 3^rd^ and 4^th^ trials than the M1–alone group (p = 0.01, 0.03). In addition, the M1–IPL group showed a trend of significantly greater strength of illusion in the last 9^th^ trial (T9) than the IPL–alone group (p = 0.055; Figure 6A). We also examined difference between the M1–IPL and IPL–IFG groups. A mixed-design ANOVA (group [2] x trials [9]) showed significant interaction (F(8, 224) = 5.59, p = 0.025, partial η² = 0.17). The M1–IPL group reported significantly greater strength of illusion in the 4^th^, 6^th^, 7^th^ and 9^th^ trials than the IPL–IFG group (p = 0.04, 0.02, 0.01, 0.02; Figure 6B). Finally, we examined difference between the M1–IPL and IFG–alone groups. A mixed-design ANOVA (group [2] x trials [9]) showed no significant interaction. The IFG–alone group showed significantly greater inter-participant variability than the M1–IPL group (SI Appendix Figure S8).

**Figure 6.**
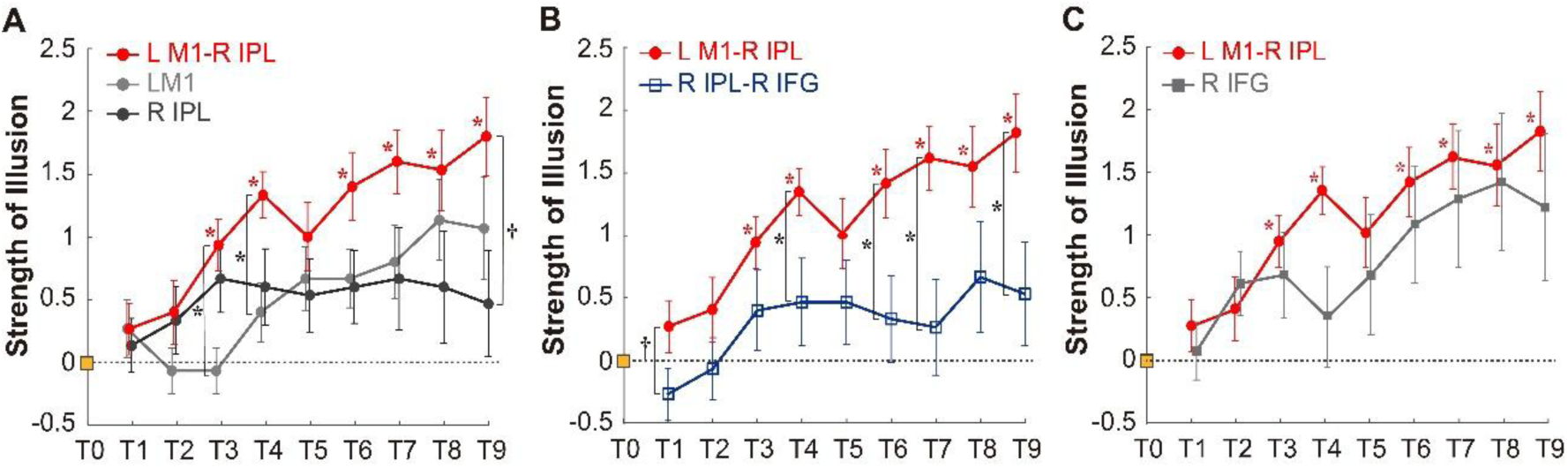
Results of the second tDCS experiment. (A) Comparison across M1–IPL, M1–alone, and IPL–alone groups. (B) Comparison between M1–IPL and IPL–IFG groups. (C) Comparison between M1–IPL and IFG–alone groups. Note that the data for M1–IPL group are identical across all panels. Red asterisks indicate significant difference when each trial (T1 to T9) was compared with baseline trial (T0) after Bonferroni correction. Black ones indicate significant between-group difference in each trial (corrected p < 0.05). † indicates a significant trend in between-group difference for each trial (corrected p < 0.1).

These findings demonstrate that the augmentation of kinesthetic illusion by M1–IPL tDCS is reproducible and robust across independent participant groups, relative to the other tDCS conditions. Together with the fMRI findings, the series of results provide converging evidence that the M1–IPL bottom-up network causally contributes to conscious kinesthetic perception.

#### Muscle activity

In the second tDCS experiment, we also measured muscle activity. The purpose of this investigation was to clarify whether or not muscle activity increases in relation to augmentation of illusion during the M1–IPL tDCS. No significant increase of muscle activity across trials was found either in the TA muscle (antagonistic muscle for the illusion), the lateral and medial Gastrocnemius muscles, or the Soleus muscle (agonistic muscles for the illusion; Figure 7 and SI Appendix). Hence, the M1–IPL tDCS augmented the illusion centrally without generating muscle activity.

**Figure 7.**
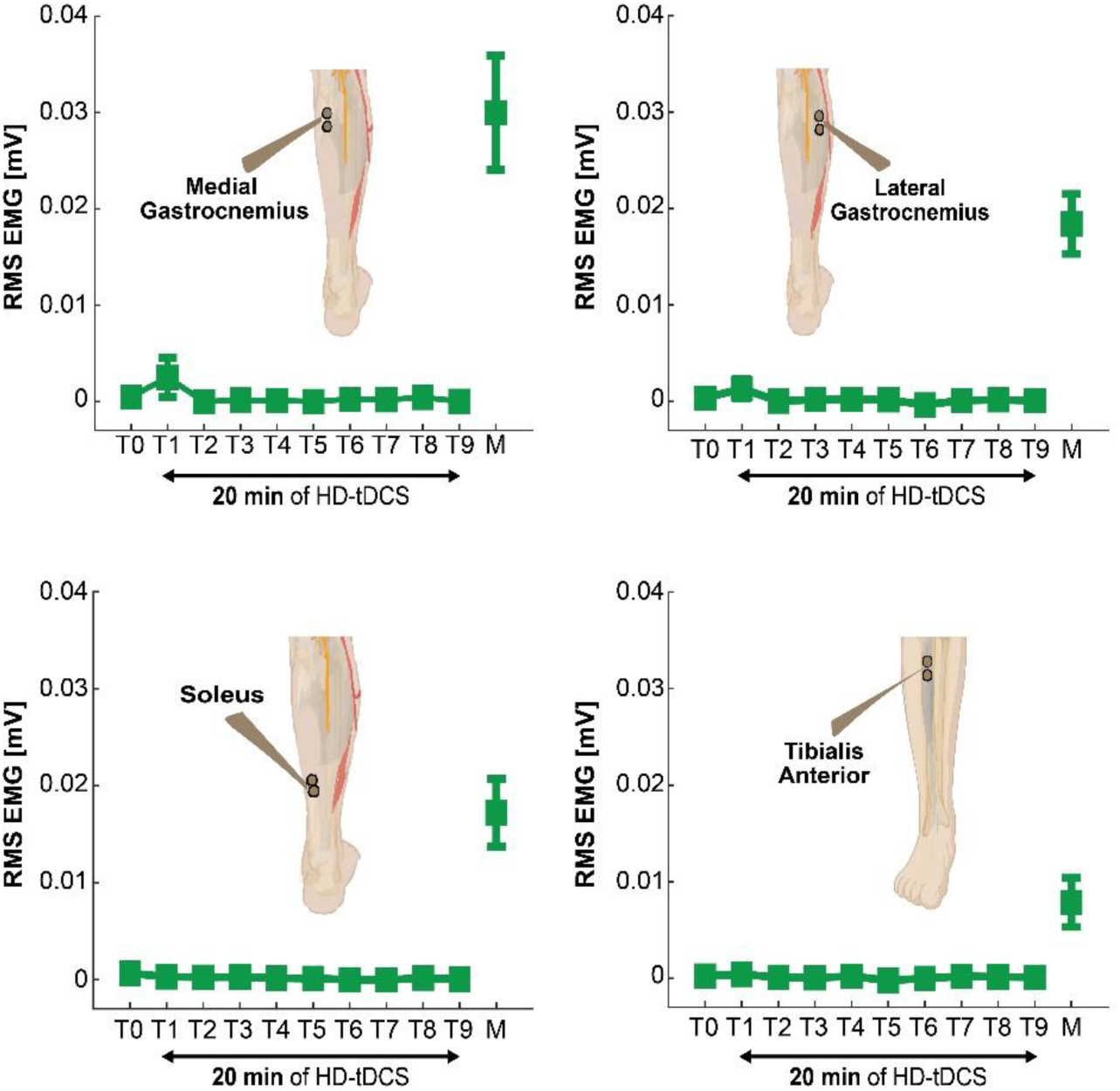
Mean EMG activity obtained from the four muscles across participants in baseline and each trial (T0-T9) during M1–IPL tDCS and during execution of plantarflexion movement of the right foot (M). We statistically evaluated if the EMG values are greater during tendon-vibration period than during resting period, and no significant differences were found for any muscle in any trial (p > 0.05). The error bars indicate the standard error of the mean across participants. In many cases, the error bars are not visible from T0 to T9 because of the small variance. EMG: electromyogram, RMS: root mean square.

## Discussion

This study provides compelling evidence that kinesthetic awareness emerges from coordinated activity between the primary kinesthetic-motor cortex (M1) and higher-order right frontoparietal cortices (IPL and IFG). The fMRI results revealed distinct but complementary roles within this network: the M1–IPL network supports bottom-up kinesthetic processing, whereas the IFG contributes to top-down monitoring (prediction) of kinesthetic experience. The tDCS results further demonstrated an essential and causal contribution of the M1–IPL network to kinesthetic awareness. Thus, conscious kinesthetic perception appears to depend not simply on activation of a modality-specific cortical region (25), but on its interaction with higher-order frontoparietal cortex, consistent with the Global Neuronal Workspace (GNW) hypothesis. Our findings extend this principle to bodily awareness, demonstrating a causal contribution of interactions between kinesthetic-motor representations in the M1 and higher-order bodily-self representations in the right IPL. This finding advances our mechanistic understanding of kinesthetic awareness and may provide insights into clinical disturbances such as anosognosia for hemiplegia.

### M1 and EMG

The M1 foot section and its adjacent SMA were activated when the participants experienced illusory plantarflexion of the stationary foot even without prior knowledge of illusion (Figure 2B). Activity in the contralateral M1 and its anteriorly adjacent area (dorsal premotor cortex for hand or SMA for foot) is most consistently observed during illusory limb movement in our series of studies (see Introduction), suggesting the primary importance of the M1 for kinesthetic illusion (12).

Kinesthetic signals from the muscle spindles may reach the M1 via the primary somatosensory area 3a and the spinocerebellum, or directly through the thalamus (8,26). In the present study, although tendon vibration was applied, M1 activity did not increase; rather, it was suppressed when the UA group experienced no illusion during the first two sessions (Figure 2C). This suppression of M1 activity appears to be consistent with previous findings showing that viewing a stationary vibrated hand attenuates the kinesthetic illusion of hand movement, with suppression of activity in the M1 hand area proportional to the degree of attenuation (see Supplemental Material in (27)). When we examined the activity of the right spinocerebellum that receives the kinesthetic inputs from the right foot (SI Appendix Figure S3), significant increase of spinocerebellar activity was found during tendon vibration (s12) even when the UA group failed to experience the illusion. This suggests that even though the right spinocerebellum likely received the kinesthetic inputs during the first two sessions in the UA group, the M1 activity did not increase (Figure 2C) and the participants did not experience the illusion. Although the underlying reason remains unclear, it is possible that the spinocerebellar activity did not sufficiently reach M1 and/or that central inhibition occurred. For example, the primary somatosensory cortex (S1), which processes vibrotactile signals, may have suppressed M1 activity (cf. 28, 29, 30). Indeed, the UA group predominantly reported vibrotactile sensation during the first two sessions. Moreover, the right spinocerebellar activity during s12 was significantly greater in the AA group than in the UA group (SI Appendix Figure S3), supporting our claim that the illusions experienced by the AA group during s12 largely depended on bottom-up kinesthetic processing.

In non-human primates, precentral (M1) neurons that respond to passive wrist flexion also respond to tendon vibration of wrist extensor muscles, which can be potential neural mechanism in the M1 underlying illusory wrist flexion (26). In humans, cortico-spinal excitability for the wrist flexor muscles increases when participants experience illusory wrist flexion during tendon vibration of wrist extensor muscles, which is the same pattern during actual wrist flexion (31). It is likely that illusory limb movement arises from a neural circuit in which M1 neurons receive Ia afferent input from a muscle and send motor commands to its antagonistic muscle (32). Importantly, robust activation of the agonistic muscle, as observed during actual movement, is normally absent during illusory movement (Figure 7; (14)), though it is also true that muscle activity occasionally emerged in the agonistic muscles during illusion in the first tDCS trial (T1) of two participants. Thus, in most cases of illusory limb movement, kinesthetic signals appear to be transformed into latent motor commands within the M1 (2). This latent neural facilitation of the agonistic (non-vibrated) muscles in the M1 likely underlies the experience of illusory limb movement, and this neural state is likely to become conscious through its dynamic interplay with the right IPL (see next).

### Right IPL and M1-IPL network

Both the right IPL and the M1, in addition to the SMA, were activated even when participants experienced the illusion without prior knowledge (Figure 2B). The right IPL enhanced its functional coupling with the M1 during illusion with knowledge (Figure 3A). These findings indicate that these regions constitute a functional network that supports bottom-up kinesthetic processing. Since direct causal influence from the M1 to the IPL was observed regardless of whether participants experienced the illusion with or without knowledge and even when the UA group failed to experience the illusion without knowledge (Figure 3B), a bottom-up flow of kinesthetic information from the M1 to the right IPL could be a fundamental feature of this network. Through this pathway, the IPL may continuously represent the neural state of the M1. For example, when participants experience the illusion, the IPL may represent the M1 state associated with facilitation of the agonist muscles (see above). It is likely that concurrent dual-site tDCS to the M1 and the right IPL may facilitate these neuronal processes and augment kinesthetic awareness (Figures 5, 6). In contrast, when participants do not experience the illusion, the IPL may become silent by reflecting the suppressed or inactive state of the M1 (Figure 2C top two panels).

Direct electrical stimulation of the right IPL can evoke illusory limb movements (33). Moreover, viewing a video of hand flexion during an illusory hand-flexion enhances both the illusion and right IPL activity, whereas viewing hand extension attenuates both (34). These findings highlight the important role of the right IPL in kinesthetic awareness. In the present study, however, enhancement of the illusion was observed with the M1–IPL tDCS, but not with the IPL–alone tDCS (Figure 6A). Rather than contradicting the role of the IPL, this finding suggests that IPL–alone tDCS may be insufficient to enhance kinesthetic awareness and that its interaction with the M1 is critical for generating or strengthening the conscious kinesthetic experience. Thus, the right IPL may serve as a higher-order integrative hub whose contribution to kinesthetic awareness depends on its interaction with modality-specific kinesthetic representations in the M1.

This M1–IPL network may support kinesthetic awareness across multiple body parts, as the right IPL likely receives and integrates kinesthetic signals from both the hands and feet through the M1 (4). Moreover, a region highly similar to the right IPL area activated during the illusion is also engaged during self-face recognition, a process closely linked to bodily self-awareness (35). This region selectively responds to one’s own face rather than others’ faces, suggesting that the right IPL may contribute not only to kinesthetic awareness but also more broadly to the representation of the bodily self.

A comparable region in the right IPL has also been shown to play a key role in proprioceptive–motor matching between the left and right feet (36) and in proprioceptive–motor coupling between the left and right hands (37). These findings suggest that the right IPL is important not only for kinesthetic awareness but also for proprioceptive–motor integration. Adjacent to the right IPL region where electrical stimulation evokes kinesthetic awareness lies another area in which stimulation elicits a conscious intention to move (33). Furthermore, transcranial magnetic stimulation over the right IPL has been reported to disrupt sense of agency (motor intention; 38, 39). Thus, just as the M1 forms a circuit linking motor commands with kinesthetic feedback, the right IPL may constitute a higher-order circuit that integrates and compares motor intention with kinesthetic awareness at the level of conscious experience (c.f. (40)). In this sense, the M1–IPL network can be considered to be involved in more general motor awareness. It is therefore plausible that concurrent dual-site tDCS over the M1 and the right IPL not only augments kinesthetic awareness (Figures 5 and 6) but also facilitates proprioceptive–motor integration, a process that is essential for motor skill acquisition and recovery of motor function after stroke (37). Future studies are needed to test these possibilities.

### Right IFG

Unlike the M1 and the right IPL, the right IFG (area 44) selectively increased its activity when UA group noticed and experienced the illusion with knowledge (Figure 2B blue section). This indicates that the IFG has a distinct role from the IPL, though these are tightly connected both functionally (16) and structurally (14, 15, 41). This region is likely involved in the top-down monitoring, expectation, and prediction of kinesthetic signals or experiences during tendon vibration when a prior knowledge of illusion is given. Direct causal influence from the IFG to the M1 and the IPL was consistently observed when the UA and AA groups experienced the illusion with knowledge, suggesting that the IFG (area 44) exerted top-down influences both to the M1 and to the IPL during illusion with knowledge (Figure 3B). The top-down influence from the IFG to the M1 was distinct from that observed when the AA group experienced the illusion in a bottom-up manner (Figure 3B). The M1 became active when the UA group became aware of the illusion with knowledge (Figure 2C top left). We may speculate that the top-down influence from the right IFG may augment the M1 activity centrally (Figure 2C) when the UA group noticed the illusion (s34) even in the situation that the right spinocerebellum did not show robust activity (SI Appendix Figure S3). This view is compatible with the previous claim that attempting to move the right foot during ischemic nerve block activates the right IFG/PMv, and this activity likely compensates for the reduced activation in the primary sensorimotor foot area by transmitting predictive signals (18).

Although the M1–IPL group demonstrated a significant enhancement of illusion strength, whereas the IFG-alone group did not, no significant between-group difference was detected (Figure 6C). This lack of significance was likely due to the markedly greater inter-participant variability observed in the IFG-alone group relative to the M1–IPL group (SI Appendix Figure S8). Therefore, the extent to which tDCS applied to the right IFG alone can elicit an augmentation effect comparable to that produced by dual-site M1–IPL tDCS remains to be determined.

### Clinical implications

This study also provides important insights into clinical symptoms caused by the right hemisphere damage. First of all, activations of the right IPL and the right IFG most likely connected by the inferior branch of the superior longitudinal fasciculus (SLF III; (14, 42)) during kinesthetic illusion with knowledge fits well to the clinical report that damage to the right frontoparietal fibers (e.g., SLF III and arcuate fasciculus) disrupts proprioception (13). The results also provide important insights into anosognosia for hemiplegia in the absence of spatial neglect following right hemisphere damage (17, 43). Our findings seem to suggest that this disease appears to arise from a combination of multiple impairments: a deficit in proprioception (kinesthesia) associated with the IPL damage; disruption in monitoring the state of the motor system responsible for kinesthetic processing due to the IFG damage; and reduced awareness to general bodily stimulation resulting from lesions in the insula and putamen (Figure 2B).

### tDCS and GNW hypothesis

Illusion-related activity identified in the tDCS experiment 1 (Figure 5A) was highly similar to that observed in the fMRI experiment (Figure 2A). In addition, increased activity in the right spinocerebellum observed across all groups in the tDCS experiment 1 (SI Appendix Figure S7) suggests that bottom-up processing also contributed to the illusion. Thus, the brain stimulation in the tDCS experiment 1 was likely applied under conditions in which both bottom-up and top-down processes contributed to the kinesthetic illusion. In the current study, we aimed to achieve dual-site, yet spatially localized, tDCS of the target regions. To this end, we adopted a novel approach involving the simultaneous stimulation of two sites using pairs of electrodes. Despite this new apporach, the present local tDCS approach appears to have effectively stimulated the left M1, as well as the right IPL and right IFG (Figures 5A, B, SI Appendix, Figures S5, S6). It is shown that tDCS may facilitate neuronal processing by augmenting synaptic transmission in a stimulated region (44). Hence, it is likely that the current tDCS applied to the M1, the right IPL and the right IFG could facilitate their local neuronal processing.

Although the M1, right IPL, and right IFG form an important network for experiencing the illusion, a robust augmentation effect was observed only with M1–IPL tDCS (Figures 5 and 6). The reason why the other tDCS conditions did not effectively enhance the illusion remains unclear. One possibility is that the effects of brain stimulation depend on complex linear and nonlinear interactions among stimulated and interconnected brain regions. For example, the effect of M1–IPL tDCS appeared to be comparable to the combined effects of M1-alone and IPL-alone tDCS (Figure 6A), suggesting a possible additive effect. In contrast, the augmentation effect differed between the M1–IPL and M1–IFG conditions despite stimulation of the same M1 region (Figure 5C). Thus, the effects of tDCS may depend not only on which region is stimulated, but also on how the stimulated regions interact within the underlying network.

Although two brain regions were stimulated simultaneously, the effects of M1–IPL tDCS were more robust than those of M1–IFG and IPL–IFG tDCS (Figures 5C and 6B). These findings suggest that the M1–IPL bottom-up network plays a more direct and causal role in kinesthetic awareness. This interpretation is consistent with the inherently bottom-up nature of kinesthetic illusion, particularly the aftereffect—the sensation that the limb returns toward its original stationary position after the vibration ceases (31)—which cannot be experienced during top-down motor imagery. Thus, bottom-up sensory processing appears to be essential for generating kinesthetic illusions.

Finally, the GNW theory proposes that conscious sensory perception emerges from dynamic interactions between modality-specific sensory cortices and distributed frontoparietal networks. Our findings provide causal evidence that coordinated interactions between the modality-specific kinesthetic-motor cortex (M1) and the higher-order right inferior parietal cortex (IPL) contribute to kinesthetic awareness, suggesting that the GNW framework may also apply to bodily awareness beyond exteroceptive perception.

### Limitations

A limitation of the present study is the limited spatial focality of tDCS. Although the bilateral SMA has been identified as a component of the bottom-up network and has been suggested to play a secondary role in integrating kinesthetic signals from multiple limbs and distributing these signals across the two hemispheres (4), the possibility that the observed effects were partly attributable to SMA stimulation cannot be completely excluded. Nevertheless, the estimated electric field (EF) strength in the bilateral SMA ROIs was lower than that in the left M1 ROI in both the M1–IPL and M1–IFG groups (SI Appendix and Figure S6C, D), suggesting that the stimulation preferentially targeted the left M1. However, the EF generated by tDCS may have extended to the adjacent SMA and other cortical regions. Moreover, the EF in the M1–IFG group appeared to extend more broadly into the right hemisphere than that in the M1–IPL group (Figure 5B; SI Appendix and Figure S6A, B), raising the possibility that differences in stimulation spread may have contributed to the observed group effects. Thus, although our EF simulations support preferential stimulation of the left M1, the inability of tDCS to provide perfectly focal stimulation represents an inherent limitation, and contributions from neighboring regions, including the SMA, cannot be entirely ruled out.

## Materials and methods

For the fMRI and tDCS experiments, the experiment protocol was approved by the Ethics Committee of the National Institute of Information and Communications Technology (NICT), and the MRI Safety Committee of the Center for Information and Neural Networks (CiNet; no. 2003260010). Before the experiments, the participants received detailed descriptions regarding the experiment and subsequently provided written informed consent. The experiment was conducted according to the principles and guidelines of the Declaration of Helsinki (1975). This study basically followed the Consolidated Standards of Reporting Trials (CONSORT) guideline (2010).

### fMRI experiment

#### Participants

Sixty-two healthy young adults (mean ± standard deviation: age, 21.2 ± 1.5 years, 28 males) participated in the fMRI experiment. We verified the absence of participants with a history of neurological, psychiatric, and motor disorders based on their self-report. We determined the dominantly operating foot by asking, “Which leg do you prefer to use to kick a ball?”. This question has been shown to be valid to determine the dominantly operating foot (45), and was used consistently in the two tDCS experiments. In the fMRI experiment, 61 of the 62 participants were right-foot dominant, and the remaining one reported bilateral footedness.

#### fMRI data acquisition

Functional images were collected using T2*-weighted two-dimensional gradient echo-planar imaging (EPI) sequences on a 3.0-Tesla MRI scanner (MAGNETON TrioTim, Siemens, Germany) with a 32-channel array coil. Each functional volume consisted of 44 slices with a slice thickness of 3.0 mm, covering the entire brain. The time interval between successive acquisitions from the same slice was 2500 ms. We used an echo time (TE) of 30 ms and flip angle (FA) of 80°. The matrix size was 64 × 64 pixels, and the voxel dimensions were 3 × 3 × 3.51 mm in the x-, y-, and z-axes, respectively. For each experimental run, we collected 65 volumes.

For an anatomical reference, we acquired a T1-weighted magnetization-prepared rapid gradient echo (MPRAGE) image using the same scanner. The imaging parameters were as follows: repetition time = 1,900 ms, TE = 2.48 ms, FA = 9°, FOV = 256 x 256 mm^2^, matrix size = 256 x 256 pixels, slice thickness = 1.0 mm, voxel size = 1 x 1 x 1 mm^3^, and 208 continuous transverse slices.

#### Experimental task

The participants rested on a bed in the supine position inside an MRI scanner. Their heads were immobilized using sponge cushions and adhesive tape to reduce head movements, and their ears were plugged for noise protection. We instructed the participants to close their eyes, relax their bodies, avoid unnecessary movements, and think of the assigned task during the fMRI scanning.

Each participant completed four sessions, each of which composed of five 15-sec tendon-vibration (Tendon) and five 15-sec bone-vibration (Bone) epochs, 150-sec in total (Figure 1A). In Tendon, we vibrated the tendon of the TA muscle of the relaxed right foot. We used the bone-vibration as a cutaneous control for the tendon-vibration task following our previous studies (35, 42), during which all participants felt vibration sensations without experiencing any reliable illusions.

Either the tendon of the muscle or the skin overlying the lateral malleolus of the right foot was vibrated using a non-magnetic vibrator (ILLUSOR, Umihira Ltd., Kyoto, Japan), driven by constant air pressure supplied by an air compressor system (see SI Appendix). This system provides a stable and appropriate vibration frequency (approximately 70 Hz) for inducing kinesthetic illusions and has been used in our previous study (46). Details of vibratory stimulation are described in SI Appendix and Figure S4. The contact surface on the skin was approximately 1 *cm*^2^ for the tendon-vibration and bone-vibration conditions. An experimenter (E.N.) applied the vibration to the skin manually with light pressure. Computer-generated auditory instructions were given to the experimenter to instruct him to start and stop Tendon and Bone epochs. The participants could not hear these instructions.

Unlike our series of previous studies, in this study, all participants did not have prior knowledge about vibration-induced illusory movement before entering the scanner. The participants closed their eyes. In the first two sessions (sessions 1 and 2, referred to as s12), we vibrated the right foot without giving them prior knowledge of vibration-induced illusory movement. After each session, the participants were asked to report any sensations they experienced in the vibrated right foot. In the last two sessions (sessions 3 and 4, referred to as s34), we vibrated the right foot after the knowledge was given. The participants were requested to be aware of movement sensations of plantarflexion of the right foot during tendon vibration and to report them after the session. After each session, when the participants reported the movement sensations, we asked them to report maximum illusory angles that they experienced. In the scanner, we showed the participants a protractor on which a foot-shape indicator was mounted. The participants viewed this picture through a mirror attached to the MRI coil in front of their eyes. This indicator was first set at the foot angle in the relaxed position of each participant. From this position, we began plantarflexing the indicator. When the participants believed that the indicator had reached the maximum illusory angle that they experienced, they were asked to say “stop.” We measured this angle as a change from the original position. We calculated mean maximum illusory angle for all participants during sessions 1 and 2 and during sessions 3 and 4 in the UA and AA group. One-sample t-test was performed to examine whether the mean illusory angle across participants is significantly different from zero (SI Appendix Figure S1).

### Functional image analysis

#### Preprocessing

To eliminate the effects of unsteady magnetization, the first four EPI (echo-planar) images were discarded in each fMRI session. We analyzed imaging data using Statistical Parametric Mapping, SPM 12 (default setting: Wellcome Trust Centre for Neuroimaging, London, UK) implemented in MATLAB R2024a (MathWorks, Sherborn, MA, USA). The following preprocessing steps were performed for each participant using default SPM parameters unless otherwise specified. The EPI images were realigned to the first image and then to the mean image to correct for head movements. Through the realignment, time-series of the head position were obtained by a rigid body transformation (linear transformation) using the least squares method for six parameters (translational displacements along the x-, y-, and z-axes and the rotational displacements of pitch, raw, and roll). We calculated the framewise displacement (i.e., the absolute value of displacement in each frame from its previous frame, FD) based on the six parameters (47). The majority of the participants had no frames in which FD exceeded 0.9 mm (60/62). Even in participants who had such frames, the percentage was less than 2% of the total volumes. Therefore, no data were excluded from the analysis. Next, the T1-weighted structural image of each participant was coregistered to the mean image of realigned EPI images using affine transformation. Then, the coregistered T1-weighted structural image was spatially normalized to the standard stereotactic Montreal Neurological Institute (MNI) space. Applying the parameter estimated in this process, individual realigned images were normalized to the MNI space. Applying the parameters that were estimated in this process, individual realigned images were normalized to the MNI space with a 2-mm isotropic voxel size using the SPM12 normalization algorithm. Finally, the spatially normalized functional images were filtered using three-dimensional Gaussian function with a full width at half maximum (FWHM) of 8 mm.

#### Single-subject analysis

After preprocessing, illusion-related activity (Tendon > Bone) was estimated for each participant using the general linear model (GLM) implemented in SPM 12 (48, 49). First, a design matrix was prepared for each participant. In the single-subject analysis, the design matrix contained a boxcar function modeling the Tendon epochs in each session, convolved with the canonical hemodynamic response function (HRF). In addition, we included six realignment parameters in the design matrix as regressors of no interest to correct for the residual motion-related variance after realignment. Also, a high-pass filter with a cutoff of 128 s was applied to remove low-frequency drifts in the BOLD signal. A contrast image showing activation (Tendon > Bone) in each session was created for each participant to be used in the following second-level group analyses (4, 7, 14).

#### Group analysis

For the second-level group analysis, we used a full factorial design with a within-subject factor (knowledge [2]: w/o s12, w s34) and a between-subject factor (group [2]: AA, UA). First, we examined illusion-related activity in the last two sessions (s34) in which both AA and UA groups experienced the illusion with knowledge (Figure 2A). Next, we compared brain activity during the first two sessions (s12) between the AA and UA groups by evaluating the contrast (AAs12 - UAs12; Figure 2B). Finally, we evaluated interaction term (UA(s34 - s12) - AA(s34 - s12); Figure 2B) to examine brain regions in which activity increased selectively when the UA group noticed and experienced the illusion while attending it, compared with when they did not, by controlling for the session order effect. In these analyses, we used a small volume correction (SVC) approach (uncorrected voxel-wise threshold of p < 0.005 and extent threshold of p < 0.05, FWE-corrected) within the predefined ROI (see SI Appendix and Table S2). We used the probabilistic cytoarchitectonic maps available in the SPM anatomy toolbox v3.0 to identify anatomical regions of activation peaks (50, 51). Finally, in order to visualize the illusion-related activity change in the UA and AA group, we extracted individual brain activity (parameter estimates) from each significant cluster (e.g., the left M1-SMA, the right IPL, the right IFG and the left insula) for each participant. Then, we calculated the mean and standard error of the mean (SEM) across participants in each group for s12 and s34 (Figure 2C).

#### Functional connectivity analysis

We conducted a generalized psychophysiological interaction (gPPI) analysis to identify brain regions in which activity enhanced functional coupling with seed regions during illusion with knowledge. In this analysis, the raw EPI images were preprocessed and analyzed using the CONN functional connectivity toolbox (version 22.a), implemented in SPM 12 (52, 53). Physiological noise originating from white matter and cerebrospinal fluid (54), head motion-related artifacts (55), and scrubbing and session effects (56) were removed. Temporal band-pass filter of 0.008–0.09 Hz was applied to examine task-related functional connectivity change within the slower range-brain activity fluctuations, and to reduce the noise related to respiratory (∼0.3 Hz) and cardiac (∼1 Hz) fluctuations (57).

For the gPPI analysis, we selected five regions as seeds based on the group results of the contrast analyses (Figure 2A): the left M1, left or right SMA, right IPL, and right IFG. The seed regions were then defined based on the predefined ROI (SI Appendix Table S2). Each seed region was defined as a sphere of 10-mm radius centered at the peak coordinates. The 10-mm radius was selected by considering final smoothness of functional images (about 9.2 mm FWHM). After nuisance regression, each participant’s time course of the average fMRI signal across the voxels in each seed region was deconvolved using a canonical hemodynamic response function (physiological variable). Then, we performed a general linear model analysis using the design matrix with the following regressors: the physiological variable, boxcar function for the task epoch (psychological variable), and multiplication of the physiological variable and the psychological variable (PPI regressor). These variables were convolved with a canonical hemodynamic response function, and six realignment parameters were also included in the design matrix as regressors of no interest.

We first generated an image of voxels showing to what extent their activities changed with the PPI regressor of each seed region in the first two and last two sessions (s12 and s34) separately for each participant. Then, in the group-level analysis, we generated a contrast image ([UA(s34 - s12)] - [AA(s34 - s12)]) to examine brain regions which enhanced functional connectivity with each seed region when the UA group noticed the illusion with the knowledge as compared to when they did not without the knowledge, by eliminating session order effect, i.e. interaction term. For this analysis, we adopted SVC approach to identify significant activity (Height threshold, p < 0.005, uncorrected; extent threshold, p < 0.05, FWE-corrected).

#### Causality analysis using the Linear Non-Gaussian Acyclic Model (LiNGAM)

We used the LiNGAM to investigate the causal relationships among the three regions (the left M1 foot section, the right IPL and the right IFG) during tendon vibration (illusion). The LiNGAM (58) is a statistical method for discovering structural causal relationships among time-series data. This allows us to explore the causal relationships (orders) across multiple brain regions without requiring prior knowledge or specific hypotheses for the network structure (59, 60). Based on observational data, it estimates directed causal relationships, which may be either positive or negative, within an unknown causal structure. The LiNGAM is expressed as follows:

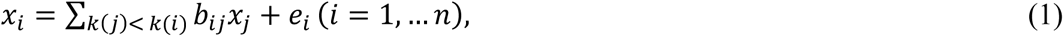

where the observed variable *x_i_* is represented as a linear combination of the other observed variables *x_j_* (*j* ≠ *i*, *j* = 1, … *n*) and external influence *e_i_* (*i* =, …, *n*). The causal ordering of variable *x_i_* is represented by *k*(*i*). The external influences *e_i_* are assumed to be an independent and non-Gaussian distribution with a zero mean and nonzero variance. This assumption is crucial for discovering the causal ordering of the variables. The connection weight *b_ij_* represents the causal effect of the observed variable *x_j_* on *x_i_*. Equation (1) can be rewritten in matrix form as follows:

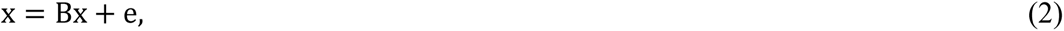

where **x** and **e** are vectors containing *x_i_* and *e_i_*, respectively. Since the LiNGAM assumes an acyclic graph structure, a causal order exists among the observed variables. Consequently, when the rows and columns are rearranged according to the causal ordering, the *n* × *n* matrix **B**, constituted by *b_ij_*, will be lower triangular.

In this study, we used DirectLiNGAM (61) to estimate the connection weights *b_ij_* and the causal ordering among the brain activities. The DirectLiNGAM does not require algorithmic parameters, such as initial values, step sizes, or convergence criteria and showed good convergence to the right solution within a small number of iterations than the original LiNGAM (61). We conducted the analysis using an open-source LiNGAM library (62). To apply the DirectLiNGAM to small-sample fMRI data, we used an approach in which data points were aggregated across participants (59, 63, 64). Figure S2 represents construction of the fMRI data matrix for the analyses (see SI Appendix).

We assessed the statistical significance of the estimated causal effects using a bootstrap method. Based on previous studies (59, 61, 64), we generated 1,000 bootstrap data matrices for each of s12 of AA, s34 of AA, s12 of UA, and s34 of UA. We then used a two-sided Wilcoxon signed-rank test to determine whether each estimated connection weight differed significantly from zero (p < 0.001). In addition, we used the Shapiro-Wilk test to confirm whether the estimated external influences e = (I − B)x satisfied the assumption of non-Gaussianity in the LiNGAM (p < 0.01).

### tDCS experiment 1

#### Study Design

This was a double-blind, parallel, randomized, sham-controlled study. The current tDCS experiments were conducted by following Japanese guidelines for the safety of low-intensity transcranial electrical stimulation (2019 Edition). Before we started tDCS experiment, we collected a low-resolution 3D structural MRI image to determine electrode positions. We also collected a T1-weighted structural image and functional MRI images in order to identify illusion-related activity and to simulate distribution and strength of electric field induced by tDCS in each participant (see SI Appendix and Figure S5 and S6).

#### Sample size

Sample size was determined using G*Power 3.1.9.7 software (65), which considered the use of a 2-factor mixed design ANOVA with one between-subject factor (3 groups) and one within-subject factor (9 trials). A medium effect size (f = 0.25) was assumed based on previous non-invasive brain stimulation studies (66, 67), with an alpha level of 0.05 and statistical power of 0.95. The computed, required total sample size was 63. We decided to collect a total of 63 participants and assign 21 participants to each group (M1–IPL, M1–IFG, and sham).

#### Electrode positioning

In the tDCS experiment, prompted by the fMRI findings, we aimed to stimulate the left M1 foot section, the right IPL, and the right IFG. We first determined electrode positions using a simulation software (HD-Explore^TM^, Soterix Medical, Inc., New York, USA). The simulation suggested that placing the anode over Cz and the cathode over C1 could provide relatively focal stimulation of the left M1 foot section. Similarly, it suggested that the anode over CP6 and the cathode over CP4 and the anode over FC6 and the cathode over FT8 could stimulate the right IPL and the right IFG, respectively. The participant wore an HD-tDCS cap (Soterix Medical, Inc., New York, USA) with electrode holders containing MRI-visible markers attached at these locations. These appear hyperintense on structural MRI images. We identified the Cz position based on the international 10–20 system in each participant and fitted the cap’s Cz position to the participant’s Cz position.

In order to check electrode (marker) positions relative to the cortex, a low-resolution 3D structure image was acquired using a 3.0-Tesla MRI scanner (MAGNETOM Vida; Siemens, Germany) equipped with a 64-channel array head-neck coil. A 3D gradient echo sequence was used (TR/TE = 3.15/1.37 ms; FA = 8°; FOV = 260 × 260 mm^2^; matrix size = 160 × 160 pixels; slice thickness = 1.6 mm; voxel size = 1.6 × 1.6 × 1.6 mm^3^; 128 continuous sagittal slices). Using this image, we checked whether the Cz and C1 markers are located over the left precentral gyrus in each individual. When the Cz position was not located over the medial (paracentral) portion of the left precentral gyrus and when the C1 position was not located around the hand knob structure of it, we adjusted these positions and scanned again to verify the positions (SI Appendix Figure S5 C). The perpendicular distance from each anodal marker to the target cortical surface was approximately 2 cm for all three regions (2.4 ± 0.3 cm at Cz, 2.11 ± 0.26 cm at CP6, and 1.99 ± 0.23 cm at FC6). No adjustment was required in 35 participants. In the remaining 28 participants, the cap was adjusted by an average of 0.68 ± 0.60 in the anterior direction and 0.3 ± 0.74 cm in the medial direction, indicating electrode positions can be determined with reasonable accuracy using the 10–20 system alone.

#### Participant selection

Given that susceptibility to illusions appears to exhibit sex differences in the fMRI experiment (SI Appendix), only male participants were recruited for the following tDCS experiments. Furthermore, to ensure a more homogeneous sample, we selected male participants classified as the UA group. We tested a total of 83 male participants with no history of neurological or other diseases. None of them had prior knowledge about vibration-induced illusory movement before the experiment. In the scanner before fMRI scanning, we vibrated the right foot for 15 seconds without giving them the knowledge. The participants were asked to report any sensations they experienced in the vibrated right foot after the vibration. This was done twice. Next, we vibrated the right foot after the knowledge was given. The participants were requested to be aware of movement sensations of plantarflexion of the right foot and to report them after the vibration. Of the 83 participants, sixty-three participants (mean ± standard deviation: age, 22.3 ± 1.7 years; 59 with right footedness, 1 with bilateral footedness, and 3 with left footedness) reported the illusion only after the knowledge was given but not before (UA group). Twenty-one participants were randomly assigned to each group (M1–IPL, M1–IFG, and sham). No significant differences in age were observed among the three groups (F(2,60) = 3.15, p = 0.55, ηp² = 0.02). Nine participants were classified as the AA group, seven participants did not report the illusion explicitly without knowledge, but reported it retrospectively, and four participants experienced no illusion. After the MRI scan, the participants wearing the HD-tDCS cap were moved to another room for the subsequent tDCS experiment.

#### Application of tDCS

The study employed a double-blind design in which both the participants and the experimenter (E.N.) were blinded to the stimulation conditions (M1–IPL, M1–IFG, and sham). Additionally, during the experiment, the stimulator display was concealed to ensure that neither the participants nor the experimenter were aware of the stimulation conditions.

We prepared three groups: M1–IPL group (simultaneous tDCS to the left M1 and the right IPL), M1–IFG group (simultaneous tDCS to the left M1 and the right IFG), and sham group (sham). A Soterix Medical MxN-5 High-Definition Transcranial Electrical Stimulator (HD-tDCS; Soterix Medical, Inc., New York, USA) was used. Ag–AgCl sintered ring HD electrodes (Soterix Medical, Inc.) were placed in HD plastic holders filled with conductive gel, which were embedded in the cap and connected to the stimulator. The reference electrode was placed at AF8. The diameter of each electrode is 12 mm. Prior to electrode application, the participant’s hair and scalp were prepared to improve skin conductivity underneath the electrodes. Electrode impedances were typically 1 to 2 kΩ and maintained below 5 kΩ throughout the tDCS experiment.

Before the tDCS experiment, each participant underwent a pre-stimulation test to assess their tolerability to the tDCS montage. Each participant in the real tDCS (M1–IPL and M1–IFG) groups received 2 mA current for each region during 20 min (including an initial 15-s ramp-up and a 15-s ramp-down at the end). The sham group received sham stimulation (initial 15-s ramp-up from 0 to 2 mA followed by a 15-s ramp-down to 0 mA both at the beginning and at the end of the session). This was done to mimic the tingling sensation experienced during the real tDCS. Here, the sham stimulation pairs (M1–IPL or M1–IFG) were randomly selected.

#### Experimental Task

In the tDCS experiment, the blindfolded participants sat on a comfortable chair, and their right legs were placed on a horizontal platform and relaxed. A total of ten trials were conducted including a baseline trial (T0) with no tDCS. Each tDCS or sham trial consists of 30-sec resting period, 15-sec tendon-vibration period, followed by 90-s reporting period (Figure 4A). We vibrated the tendon of the TA muscle of the right relaxed foot (Figure 4A). We used an electro-magnetic vibrator (80 Hz, amplitude ± 0.2 mm, SL-0105, Asahi Seisakusho, Tokyo, Japan) that is controlled by an amplifier (APD-050FCA, No. B16393, Asahi Seisakusho, Tokyo, Japan). The vibrated area was about 1 *cm*^2^. An experienced experimenter (E.N.) manually operated the vibrator, applying it to the skin with light pressure.

Computer-generated auditory instructions were given to the participants and the experimenter; “3, 2, 1, start” to inform the start of each trial and “stop” for the end of each trial. The participants were requested to be aware of illusory plantarflexion of the right foot during the vibration. After the baseline trial, we confirmed whether or not they experienced the illusion. We also asked them to remember the strength of the illusion. In the subsequent tDCS and sham trials, they were asked to report whether illusion experienced in a current trial becomes stronger, approximately the same or weaker than that experienced in one previous trial during the reporting period.

#### Statistical analysis

Illusion strength during the baseline trial (T0) was defined as zero. When the participants reported that the illusion became stronger than in the one previous trial, a score of +1 was assigned; when they reported that it was approximately the same, a score of 0 was assigned; and when they reported that it was weaker, a score of −1 was given. The scores were then summed across participants to evaluate changes in illusion strength (Figure 5 and 6).

Normality of the data was confirmed using the Shapiro–Wilk test. We first evaluated whether illusion strength was significantly different from zero (T0) in each trial (T1 to T9). One-sample t-test with Bonferroni correction was performed in each group. Next, a two-way mixed-design analysis of variance (ANOVA) was performed to examine the effects of group and trial on illusion strength. Statistical significance was set at p < 0.05. Effect sizes were reported in addition to statistical significance, with partial eta squared (η2) for ANOVA results. When a significant interaction was observed, post hoc comparisons with Bonferroni correction were performed. One-way ANOVA with Bonferroni correction was performed to evaluate between-group differences for each trial.

### tDCS experiment 2

#### Study Design

This was also a double-blind, parallel, randomized study. The same procedures and analyses as those used in the first tDCS experiment were followed, with the exceptions described below.

#### Sample size

In the second tDCS experiment, we prepared five groups (M1–IPL, M1–alone, IPL–alone, IPL–IFG, and IFG–alone). The G*Power 3.1.9.7 software suggested that the computed, required sample size was 75 (medium effect size of 0.25, a power of 0.90, and α = 0.05) for a 2-factor mixed design ANOVA with one between-subject factor (5 groups) and one within-subject factor (9 trials). We decided to collect a total of 75 participants and assign 15 participants to each group.

#### Electrode positioning

The first tDCS study demonstrated that electrode positions can be determined with reasonable accuracy (within approximately 1 cm) using the 10–20 system alone, and the target regions appeared to be effectively stimulated (Figure 5B, SI Appendix and Figures S5, S6). Based on these results, electrode positions for the second tDCS experiment were determined using the 10–20 system alone.

#### Participant selection

As in the first tDCS experiment, we selected male participants classified as the UA group. We tested a total of 99 male participants naïve to the illusion with no history of neurological or other diseases. For selection, we did the same procedure in the first tDCS experiment but using the electro-magnetic vibrator in the normal experimental room. Of the 99 participants, seventy-five participants (mean ± standard deviation: age, 22.1 ± 1.9 years; 70 with right footedness, 2 with bilateral footedness, and 3 with left footedness) reported the illusion only after the knowledge was given but not before (UA group). Fifteen participants were randomly assigned to each group (M1–IPL, M1–alone, IPL–alone, IPL–IFG, and IFG–alone). No significant differences in age were observed among the five groups (F (4,70) = 2.50, p = 0.28, ηp² = 0.07). Twelve participants were classified as the AA group, and 12 participants experienced no illusion. The tDCS procedure, experimental task and statistical analysis were identical to those used in the first tDCS experiment.

#### EMG recording and analysis

In the second tDCS experiment, we recorded electromyographic (EMG) activity using surface electrodes from the medial and lateral heads of the gastrocnemius, soleus and tibialis anterior muscle of the right leg during each trial. Details of measurement and of analysis are described in SI Appendix.

## Supporting information

Supplementary Infomation

## Acknowledgments

The funding sources were not involved in the study design, collection, analysis, interpretation of data, writing of the report, or decision to submit the article for publication. The authors are grateful to CiNet MRI staff, Mr. Koji Takashima, and Ms. Mayumi Irikawa for their technical supports, and Ms. Keiko Ueyama for the illustration.

## Author Contributions

All authors contributed to conceptualization, validation and writing, review editing, and investigation. OS, TM, and JP: formal analysis; TM, JP, and EN: funding acquisition. OS and TM: visualization. OS, TM, and EN: writing, original draft preparation. TM and EN: project administration and supervision. All authors contributed to the article and approved the submitted version.

## Data Availability Statement

Datasets are available on request: The raw data supporting the conclusions of this article will be made available by the corresponding author (EN).

