## Supplementary Infomation for "Causal contribution of motor-parietal network to conscious kinesthetic perception"

**This PDF file includes:**

Supporting Information text
Figures S1 to S8
Tables S1 to S5
SI References

**Supporting Information text**

### fMRI experiment

#### Defining region-of-interest (ROI)

We defined the region-of-interest (ROI) based on the data obtained from an independent experiment in which another group of 57 young adults experienced illusory plantarflexion of the right foot with its prior knowledge. This study was originally conducted for a different purpose. In this experiment, we used the same 3.0-Tesla MRI scanner (MAGNETON TrioTim, Siemens, Germany). The participants completed 2 sessions and each session composed of six 15-sec tendon-vibration (Tendon) and six 15-sec bone-vibration (Bone) epochs, 180-sec in total. Experimental procedures were the same in the fMRI experiment (see main text). After the preprocessing and single-subject analysis described in the main text, the second-level group analysis was performed to identify illusion-related activity (Tendon > Bone; uncorrected voxel-wise threshold of $p < 0.005$ and extent threshold of $p < 0.05$, FWE-corrected). The illusion-related activity was identified in the foot section of the left primary motor cortex (M1) extending into bilateral supplementary motor area (SMA) and pre-SMA, in the bilateral inferior frontal gyri (IFG) extending into insular cortices and putamen, in the right inferior parietal lobule (IPL) and in the right anterior cerebellar paravermis (Table S2). This result is consistent with the findings of our previous studies, we therefore used this cluster image as ROI to identify illusion-related activity in the current work.

**Sexual difference between AA and UA groups**

The majority of participants in the AA group were women (20 of 22), whereas the majority of participants in the UA group were men (20 of 30). The sex distribution differed significantly between the two groups ($\chi^2(1) = 17.2$, $p < 0.001$). This observation raises the possibility of a sex difference in the tendency to experience the foot illusion in a bottom-up manner. At the peripheral level, tendon vibration is transmitted through the tendon to activate the muscle spindles in the vibrated muscle, and this proprioceptive information is conveyed primarily by Ia afferent fibers to the brain (1). Previous studies have shown that women generally have thinner tendons and muscles than men (cf. 2, 3). Consequently, tendon vibration may produce relatively greater stimulation of muscle spindles in women, potentially resulting in stronger proprioceptive afferent input to the brain. This interpretation is consistent with the greater spinocerebellar activity observed in the AA group as compared to the UA group during the first two sessions (see SI Appendix Figure S3 below). Therefore, peripheral structural differences may contribute to the observed sex difference in the experience of the illusion without knowledge. Further research is needed to determine the relative contributions of peripheral and central mechanisms to the generation of the illusion in a bottom-up manner.

**Illusory angle**

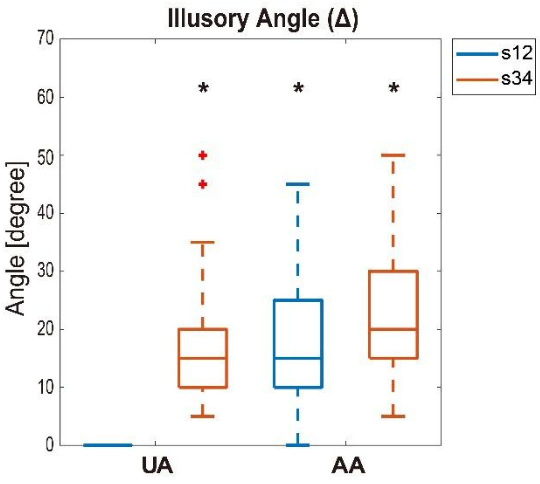


**Fig. S1.** Maximum illusory angle reported during sessions 1 and 2 (s12) and during sessions 3 and 4 (s34) in UA and AA groups. A one-sample t-test showed that the mean illusory angle across participants was significantly different from zero; $t(59) = 14.76$, $p < 0.001$, 95% CI [15.49, 20.35] for sessions 3 and 4 (s34) of UA group, $t(43) = 11.59$, $p < 0.001$, 95% CI [15.49, 22.01] for sessions 1 and 2 (s12) of AA group, $t(43) = 13.94$, $p < 0.001$, 95% CI = [18.47, 24.71] for sessions 3 and 4 (s34) of AA group. Horizontal lines within the boxes indicate the median value, while the lower and upper boundaries of the boxes represent the 25th and 75th percentiles, respectively, defining the interquartile range (IQR). Vertical dashed lines indicate the whiskers, which extend to the most extreme data points within 1.5 × IQR of the lower and upper quartiles. Data points beyond this range (outliers) are shown as red crosses. Black asterisks indicate significant difference from zero. CI: Confidence interval.

#### fMRI data matrix for LiNGAM analysis

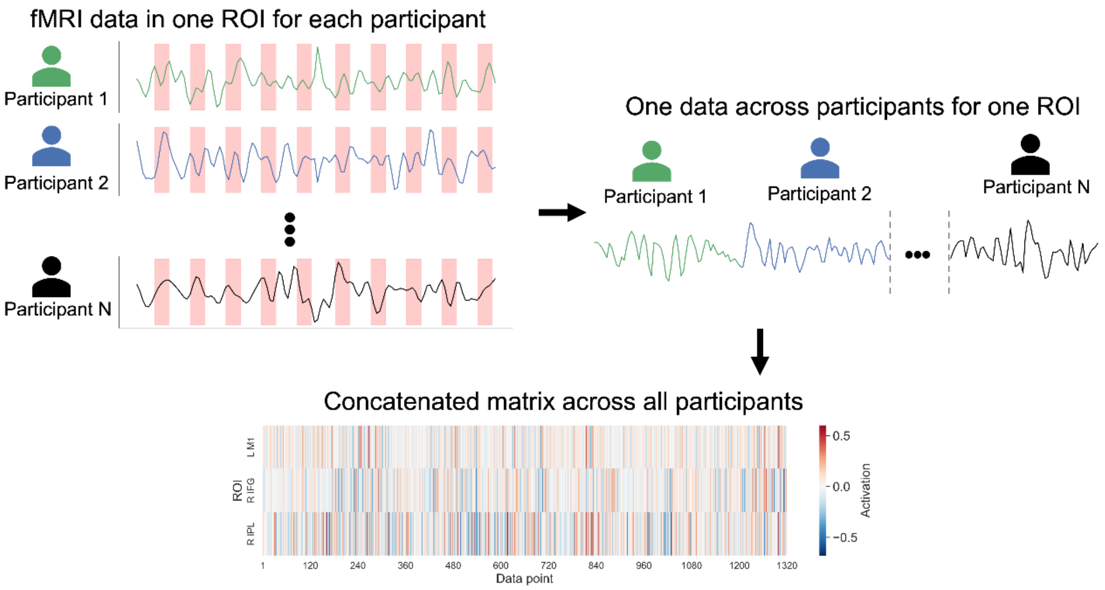


**Fig. S2.** Construction of fMRI data matrix for LiNGAM analysis. For each ROI, fMRI time-series data corresponding to Tendon epochs (red sections in top left panel) were extracted from each participant and concatenated across participants to construct the data matrix.

**Spinocerebellar activity**

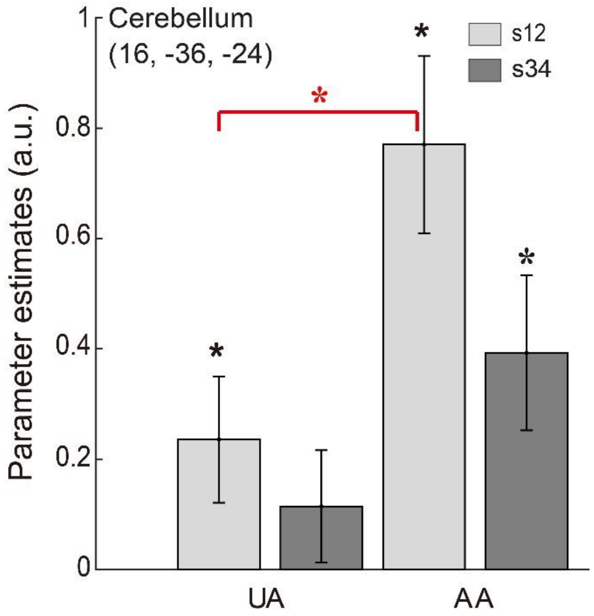


**Fig. S3.** Spinocerebellar activity in UA and AA groups. When both AA and UA groups reported the illusion during the last two sessions (s34), illusion-related activity (8 voxels) was identified in the right anterior cerebellar paravermis (peak coordinates x, y, z = 16, -36, -24) within the predefined ROI (Table S2), though this activation did not reach statistical significance. This region corresponds to the foot section of the spinocerebellum (4), which likely receives the kinesthetic (muscle spindle) signals from the right foot. We extracted individual brain activity (parameter estimates) from the cluster in each participant. We then calculated the mean and standard error of the mean (SEM) across participants in each group for s12 and s34. A one-sample t-test revealed that the mean cerebellar activity across participants was significantly different from zero ($p = 0.049$ for s12 of UA group, $p < 0.001$ for s12 of AA group, and $p = 0.011$ for s34 of AA group). The reduction in activity observed in s34 compared with s12 in both groups may be attributable to an MRI adaptation effect. Two-sample t-test showed that the cerebellar activity during s12 was significantly greater in AA group than in UA group ($t(50) = -2.79$, $p = 0.007$). Error bars indicate SEM across participants. Black asterisks indicate significant difference from zero. Red asterisk indicates significant difference between UA and AA groups in s12.

**Table S1.** Significant activations within the region-of-interest during kinesthetic illusion with knowledge (s34).

| Size | t-value | x | y | z | Anatomical identification |
|---|---|---|---|---|---|
| **2336** | 10.37 | -32 | 18 | 8 | Area Id7 |
| | 9.05 | -26 | 0 | 10 | Left Putamen |
| | 6.19 | -42 | 0 | 10 | Central Opercular Cortex |
| | 4.79 | -52 | 8 | 20 | Area 44 |
| **2449** | 7.9 | 26 | 6 | 4 | Right Putamen |
| | 7.61 | 32 | 22 | 2 | Area Id7 |
| | 7.33 | 54 | 8 | 12 | Area 44 |
| | 7.02 | 38 | 0 | 14 | Central Opercular Cortex |
| **1072** | 7.42 | -4 | -28 | 70 | Area 4a |
| | 5.69 | -6 | -12 | 66 | Area 6mc/SMA |
| | 5.64 | 6 | 0 | 58 | Area 6mr/preSMA |
| | 5.27 | -4 | -6 | 46 | Cingulate Gyrus |
| **214** | 4.54 | 62 | -20 | 30 | Area PFt/PFop (IPL) |
| | 4.33 | 62 | -24 | 46 | Area PF/PFt (IPL) |

Height threshold, $p < 0.005$, uncorrected; extent threshold, $p < 0.05$, FWE-corrected within the image showing the right-foot illusion related-activity of the 57 young adults, using SVC. Size refers to the number of significant voxels. For anatomical peak identification, we considered cytoarchitectonic areas available in the anatomy toolbox with > 30% probability (5,6). We reported the cytoarchitectonic area with the highest probability for each peak. When cytoarchitectonic areas with > 30% probability were unavailable, we simply provided the anatomical location of the peak. To facilitate visualization, we avoided reporting a peak for each cluster when it was identified in the cytoarchitectonic area or anatomical structure already reported for a peak with a higher t-value. FWE, family-wise error rate; SVC, small volume correction; IPL, inferior parietal lobule.

**Table S2.** Brain regions showing the right-foot illusion related-activity (Tendon > Bone) in an independent group of 57 healthy participants, which were used as an ROI.

| Size | t-value | x | y | z | Anatomical identification |
|---|---|---|---|---|---|
| **3163** | 10.77 | -32 | 22 | 6 | Area Id7 |
| | 9.30 | -28 | 0 | 10 | Left Putamen |
| | 8.75 | -44 | 0 | 4 | Insular Cortex |
| | 4.50 | -60 | 14 | 16 | Area 44 |
| **3364** | 8.85 | 44 | 16 | 4 | Area OP8 |
| | 8.53 | 34 | 18 | 6 | Area OP8/Id7 |
| | 7.99 | 24 | 6 | 8 | Right Putamen |
| | 7.95 | 38 | 2 | 12 | Central Opercular |
| | 6.00 | 60 | 16 | 22 | Area 44 |
| **2248** | 8.03 | -4 | -22 | 70 | Area 4a |
| | 5.88 | 10 | -2 | 66 | Area 6d2 |
| | 5.71 | -8 | -8 | 68 | Area 6d1 |
| | 5.83 | -8 | -4 | 44 | Cingulate |
| | 5.30 | 6 | 12 | 54 | Area 6mr/preSMA |
| | 3.80 | -8 | 8 | 54 | Area 6mr/preSMA |
| **496** | 5.92 | 64 | -20 | 38 | Area PF/PFt (IPL) |
| **139** | 5.04 | 14 | -34 | -30 | Paravermis (Foot Section) |
| **145** | 4.45 | 32 | -2 | 50 | Middle Frontal |

**Table S3.** Brain regions active when AA group experienced the illusion without knowledge in a bottom-up manner compared with when UA group did not [AAs12-UAs12].

| Size | t-value | x | y | z | Anatomical identification |
|---|---|---|---|---|---|
| **308** | 5.62 | -6 | -12 | 76 | Area 6d1 |
| | 5.24 | -8 | -24 | 66 | Area 4a |
| | 3.31 | -12 | -2 | 76 | Area 6d2 |
| | 3.24 | -6 | -8 | 60 | Area 6mc/SMA |
| **321** | 4.88 | 12 | 0 | 64 | Area 6d2 |
| | 4.76 | 4 | 0 | 48 | Area 6mr/preSMA |
| **166** | 4.23 | 58 | -16 | 34 | Area PFt (IPL) |
| | 4.14 | 58 | -18 | 40 | Area 2 |
| | 2.87 | 56 | -30 | 28 | Area PFcm/PF (IPL) |
| | 2.77 | 62 | -18 | 24 | PFop (IPL) |

Height threshold, $p < 0.005$, uncorrected; extent threshold, $p < 0.05$, FWE-corrected within the ROI (Table S2), using SVC. Size refers to the number of significant voxels. For anatomical peak identification, we considered cytoarchitectonic areas available in the anatomy toolbox with > 30% probability. We reported the cytoarchitectonic area with the highest probability for each peak. When cytoarchitectonic areas with > 30% probability were unavailable, we simply provided the anatomical location of the peak. To facilitate visualization, we avoided reporting a peak for each cluster when it was identified in the cytoarchitectonic area or anatomical structure already reported for a peak with a higher t-value.

**Table S4.** Right IFG region selectively active when UA group noticed and experienced the illusion while attending it, compared with when they did not, by controlling for the session order effect [UA(s34-s12)-AA(s34-s12)].

| Size | t-value | x | y | z | Anatomical identification |
|---|---|---|---|---|---|
| **136** | 3.47 | 54 | 14 | 10 | Area 44 |

Height threshold, $p < 0.005$, uncorrected; $p = 0.05$, FWE-corrected within the ROI (Table S2), using SVC. Size refers to the number of significant voxels. For anatomical peak identification, we considered cytoarchitectonic areas available in the anatomy toolbox with > 30% probability. We reported the cytoarchitectonic area with the highest probability for each peak. When cytoarchitectonic areas with > 30% probability were unavailable, we simply provided the anatomical location of the peak. To facilitate visualization, we avoided reporting a peak for each cluster when it was identified in the cytoarchitectonic area or anatomical structure already reported for a peak with a higher t-value.

**Table S5.** Functional connectivity results [UA(s34-s12)-AA(s34-s12)]

| Seed | Cluster | Size | t-value | x | y | z | Anatomical identification |
|---|---|---|---|---|---|---|---|
| **L M1** | **IFG** | **127** | 4.47 | 54 | 16 | 10 | Area 44 |
| | **IPL** | **131** | 4.15 | 60 | -30 | 40 | Area PF (IPL) |
| | | | 4.14 | 56 | -30 | 28 | Area PFcm (IPL) |

Height threshold, $p < 0.005$, uncorrected; extent threshold, $p < 0.05$, FWE-corrected within the ROI (Table S2), using SVC. Size refers to the number of significant voxels. For anatomical peak identification, we considered cytoarchitectonic areas available in the anatomy toolbox with > 30% probability. We reported the cytoarchitectonic area with the highest probability for each peak. When cytoarchitectonic areas with > 30% probability were unavailable, we simply provided the anatomical location of the peak. To facilitate visualization, we avoided reporting a peak for each cluster when it was identified in the cytoarchitectonic area or anatomical structure already reported for a peak with a higher t-value. IFG, inferior frontal gyrus.

**Evaluation of vibration frequency in MRI scanner**

In the fMRI experiment, a non-magnetic vibrator (ILLUSOR, Umihira Ltd, Kyoto, Japan) was driven by air pressure supplied by an air-compressor system. The system consists of two air compressors (Medical Compressor, Hitachi Industrial Equipment Systems Co., Ltd., Tokyo, Japan), a 1,500-L air tank (AIR TANK, ORION Thermal Engineering Co., Ltd., Nagano, Japan), and two air dryers (IDU8E-20-C-X460, SMC Corporation, Tokyo, Japan), and provides a constant air pressure of 0.75 MPa. We evaluated change of frequency of vibratory stimulation delivered by this system inside the 3-T MRI during fMRI phantom scanning. The vibrator was placed around the position where the participant's right feet are located when lying in the scanner. We operated the vibrator in the air without it coming into contact with anything. A 3D accelerometer (Acceleration Sensor MR, Brain Products GmbH, Gilching, Germany) was mounted on the contact tip of the vibrator. The three-axis accelerometer data were recorded during 160-s vibration, at a sampling rate of 5000 Hz using the Brain Products recording system (BrainAmp, Brain Products GmbH, Germany), composed of a BrainAMP EXG MR 16 channel amplifier, an ExG AUX device, and BrainVision Recorder and Analyzer softwares. The measurement was done for four times (trials). In the analysis, the data obtained from the initial 10 seconds were excluded in each trial, as we did in the fMRI analysis. First, the accelerometer signals for 150 seconds were high-pass filtered using a 4th-order Butterworth high-pass filter with a cutoff frequency of 0.5 Hz to eliminate low-frequency baseline drift and unknown DC offsets. The 150-s data were divided into 5 blocks (30-sec for each), and the peak frequency for x-, y-, z-axes was separately identified for each block using fast Fourier transform (FFT) analysis. In each block, we calculated the mean peak frequency and the standard error of the mean (SEM) across trials. The results are shown in Fig. S4. The mean peak frequencies were approximately 70 Hz for z-axis (= main direction of vibration), which are appropriate to induce kinesthetic illusion. Similar values were obtained for x- and y-axes. We also measured amplitude of vibratory stimulation using a VIPER system (POLHEMUS, Colchester, USA) with a Micro Sensor 1.8. The peak-to-peak amplitude was approximately 2.5 mm. Our previous study also used this system and confirmed that this system can elicit kinesthetic illusion (see methods in main text).

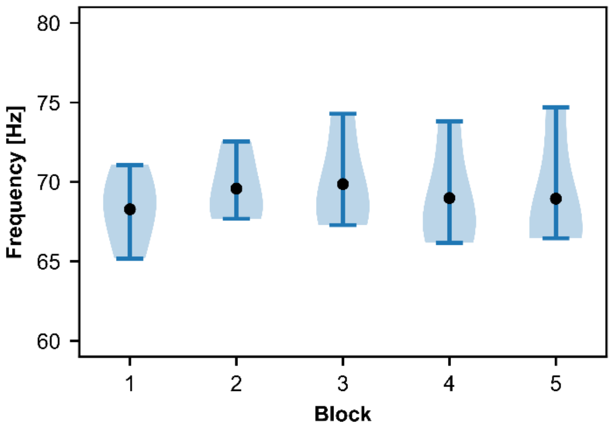


**Fig. S4.** Evaluation of vibration frequency in MRI scanner. In each block, filled circle indicates the mean peak frequency, an error bar represents the SEM, and an area shaded in light blue represents the kernel density estimate distribution of frequencies in each block.

### tDCS experiment 1

#### Identifying individual illusion-related activity

##### Data acquisition

Following acquisition of the low-resolution 3D structural MRI image (see main text), field-map images were acquired to correct for geometric distortions using a gradient-echo sequence (TR = 1,000 ms; TE1/TE2 = 5.29/7.75 ms; flip angle = 60°; FOV = 210 × 210 mm²; matrix = 70 × 70; 51 slices, voxel size = 3 × 3 × 3 mm³) in order to correct for EPI distortions caused by static magnetic field inhomogeneities. Functional images were then acquired using a T2*-weighted gradient echo-planar imaging (EPI) sequence (TR/TE = 1,000/30 ms; FA = 60°; FOV = 210 x 210 mm²; matrix = 70 x 70; 51 slices; voxel size = 3 × 3 × 3 mm³; multiband factor, 3). When obtaining functional images, each participant completed one session, consisting of six 15-sec tendon-vibration (Tendon) epochs and six 15-sec bone-vibration (Bone) epochs, with 10-sec additional resting period before the first Bone epoch, resulting in a total duration of 190-sec. In this experiment, the non-magnetic vibrator (ILLUSOR, Umihira Ltd, Kyoto, Japan) was driven by air pressure supplied by an air-compressor (EARTH MAN ACP-39SLB, Takagi, Japan). Other experimental procedures were identical to those used in sessions 3 and 4 of the fMRI experiment (see main text), in which participants received the vibration stimuli with knowledge that the vibration may induce an illusory movement. After the experiment, the EPI images were analyzed to identify illusion-related activity in each participant. A high-resolution T1-weighted MP-RAGE image was also acquired for spatial normalization (TR = 1,900 ms, TE = 2.48 ms, flip angle = 9°, FOV = 256 × 256 mm², matrix = 256 × 256, voxel size = 1 × 1 × 1 mm³, 240 contiguous sagittal slices). The individual T1-weighted images, which included MRI-visible markers for identifying the electrode positions, were also used for individualized modeling of the electric field induced by the tDCS (see below).

##### Analysis

To eliminate the effects of unsteady magnetization, the first ten EPI images were excluded from the analysis. The preprocessing procedures were almost identical to those used in the fMRI experiment (see main text), except for EPI distortion correction and the size of the smoothing kernel. For EPI distortion correction, a voxel-displacement map (VDM) calculated from the acquired field map images and used to unwarp the realigned EPI images, thereby correcting geometric distortions in the images. For spatial smoothing, a three-dimensional Gaussian kernel with a full width at half maximum (FWHM) of 6 mm was applied. In addition, framewise displacement (FD) was calculated to assess head motion. As no participant exhibited FD values exceeding 0.9 mm, no participant was excluded from the analysis on the basis of head motion.

Single-subject analyses were performed using the same procedures as those employed in the fMRI experiment (see main text). Then, a group-level one-sample *t*-test was performed to identify brain regions showing illusion-related activity across participants (Fig. S5 panel F). Based on the group-level results, peak coordinates of activation were identified for the left M1 (-4, -30, 68), right IPL (40, -40, 44), right IFG (54, 14, 16), left SMA (-8, -8, 62), and right SMA (6, -6, 66). For each region, a spherical ROI with a 10-mm radius was first defined around the group-level peak. Within each ROI, we then identified the peak of activation for each participant. When no peak was identified for a given participant, the group-level peak was used instead. Subsequently, a new spherical ROI with a 10-mm radius was centered at each participant's peak. The mean estimated electric field (EF) strength was then calculated within each participant-specific ROI for each of the five regions (see below).

**Blinding efficacy assessment**

To assess the efficacy of blinding, 45 of 63 participants were asked after the experiment: "In this brain stimulation experiment, one group received stimulation throughout the session, whereas in the other group the stimulation intensity was reduced after the initial period. Which group do you think you belonged to?" In the M1–IPL group, 11 of 12 participants (91.7%) reported that they believed they had been assigned to the group in which stimulation intensity was reduced, which was significantly greater than chance (50%; exact binomial test, $p = 0.006$). Similarly, in the M1–IFG group, 18 of 21 participants (85.7%) reported that they believed they had been assigned to the group in which stimulation intensity was reduced, which was significantly greater than chance (50%; exact binomial test, $p = 0.001$). Thus, only 4 of 33 (12 %) participants could correctly identify the real stimulation. In the sham group, 11 of 12 participants (91.7%) believed they had been assigned to the group in which stimulation intensity was reduced (exact binomial test, $p = 0.006$). Overall, the participants could not identify the stimulation they received above chance level (50%), with a correct identification rate of 33%, indicating that the blinding was effective.

**Modelling of electric field (EF) and evaluation of mean estimated EF strength in ROIs**

To visualize electric field (EF) distribution in the cortex generated by the current tDCS in each participant, EF modeling was done using SimNIBS version 4.5.0, a comprehensive open-source software that allows the simulation of NIBS-induced EFs (7). MATLAB R2024a was used to run the simulations, and Gmsh for the visualization of the results.

First, a 2 × 2 high-definition transcranial direct current stimulation (HD-tDCS) montage was modeled on the MNI-152 standard head model to estimate the EF distribution for the left M1-right IPL and left M1-right IFG stimulation. The electrode locations were determined using the 10–20 international system (Figure 4B; Cz: anode and C1: cathode for the left M1, CP6: anode and CP4: cathode for the right IPL, FC6: anode and FT8: cathode for the right IFG, and reference: AF8). Next, individual head models were generated from the individual T1-weighted structural MRI (see above) using charm (8), a free SimNIBS module that allows automated tissue segmentation from MR images, for the individualized EF simulations. The markers were visible on the T1 images, and the exact center coordinates of the markers were identified using the CONN toolbox implemented in SPM 12.

The electrodes were modeled as cylindrical structures of 12 mm in diameter and 2 mm in thickness, situated on an 8 mm gel layer with a stimulation intensity of 2 mA at the anode and -2 mA at the cathode. Simulations were performed using standard conductivity values as provided by SimNIBS (white matter: 0.126 S/m, gray matter: 0.275 S/m, CSF: 1.654 S/m, bone: 0.01 S/m, scalp: 0.465 S/m, eyeballs: 0.5 S/m, compact bone: 0.008 S/m, spongy bone: 0.025 S/m, blood: 0.6 S/m and muscle: 0.16 S/m).

The EF distribution was estimated using this simulation in the brain of each participant (Fig. S5A, B). We also calculated the mean estimated EF strength in each ROI of each participant (see above). As a reference, we also calculated the mean estimated EF strength within a sphere with a 10-mm radius centered on coordinates z, y, z = 0, -90, 2 in the non-stimulated primary visual cortex (V1). The results are summarized in Fig. S6.

Mean estimated EF strength in M1–IPL group

The mean estimated EF strength values in the left M1 and right IPL ROIs were comparable to those estimated in a previous tDCS study (9). These values were significantly higher than that estimated in the non-stimulated visual ROI ($p < 0.001$ for both after Bonferroni correction; Fig. S6A), suggesting that the left M1 and right IPL were substantially stimulated in the M1–IPL group. The mean value in the right IPL ROI was significantly higher than that obtained from the right IFG ROI ($p < 0.001$ after Bonferroni correction). On the other hand, the mean value in the right IFG ROI was similar to but significantly higher than that in the non-stimulated visual ROI ($p = 0.0015$ after Bonferroni correction, Fig. S6A). These lines of evidence suggest that the right IPL was preferentially stimulated by the tDCS targeting the IPL, though the IPL tDCS may have extended to the IFG. We also examined correlation between the mean estimated EF strength value and the augmentation effect of illusion across participants. The latter was defined as coefficient of liner regression fitted to individual data (from T1 to T9). Neither the left M1 value, the right IPL value nor the sum of the left M1 and right IPL values was significantly correlated with the augmentation effect.

Mean estimated EF strength in M1–IFG group

The mean estimated EF strength values in the left M1 and right IFG ROIs were significantly higher than that estimated in the non-stimulated visual ROI (both $p < 0.001$ after Bonferroni correction; Fig. S6B), suggesting that these regions were substantially stimulated in the M1–IFG group. The mean value in the right IFG ROI was significantly higher than that obtained from the right IPL ROI ($p < 0.001$ after Bonferroni correction). The mean value in the right IPL ROI was also significantly higher than that estimated in the non-stimulated visual ROI ($p < 0.001$ after Bonferroni correction, Fig. S6B). These lines of evidence suggest that the right IFG was preferentially stimulated by the tDCS targeting the IFG, though the IFG tDCS may have extended to the IPL. Neither the left M1 value, the right IFG value nor the sum of the left M1 and right IFG values was significantly correlated with the augmentation effect across participants.

The EF in the M1–IFG group appeared to extend more broadly into the right hemisphere than that in the M1–IPL group (Figures 5B and S5A, B). The stronger EF in the right IFG (Fig. S6B) might be related to the shorter distance between the electrodes and the cortical surface for the right IFG tDCS ($1.99 \pm 0.23$ cm at FC6) than for the right IPL tDCS ($2.11 \pm 0.26$ cm at CP6, $p < 0.05$). This stronger EF in the right IFG might extend into the right IPL.

Extension of M1-targeting tDCS to SMA

M1-targeting tDCS may have extended to the adjacent SMA (Fig. S6C, D). However, in both the M1–IPL group (Fig. S6C) and the M1–IFG group (Fig. S6D), the mean estimated EF strength was significantly lower in the left and right SMA ROIs than in the left M1 ROI. These findings indicate that M1-targeting tDCS preferentially stimulated the left M1, although the EF generated by the tDCS may have extended to the adjacent SMA.

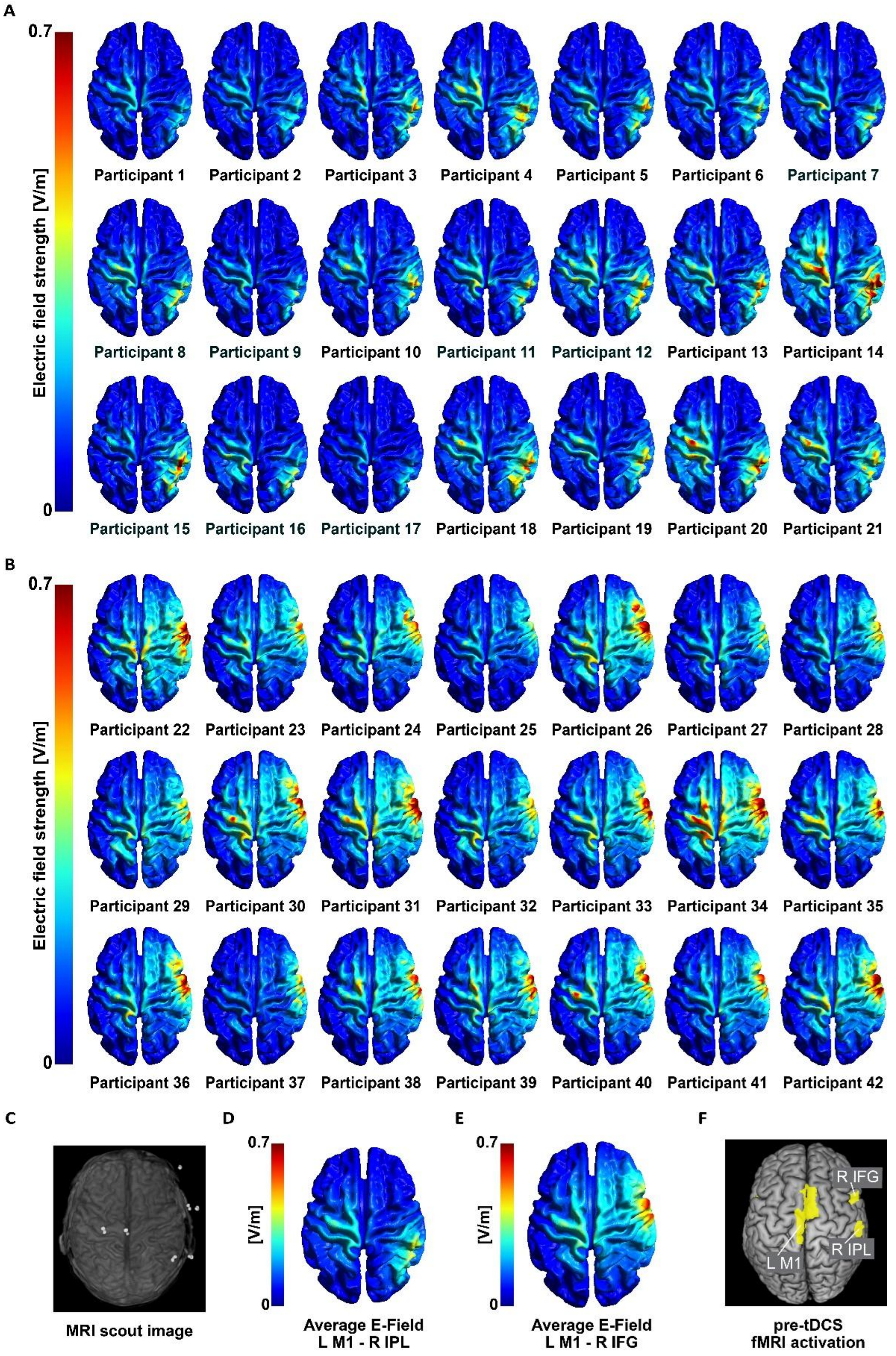


**Fig. S5.** Simulation of tDCS. (A), (B) Distribution of electric field (EF) in each participant of the M1–IPL (A) and M1–FG (B) groups. The EF strength is visualized using a color scale ranging from blue to red. (C) A typical example of a low-resolution 3D structure image including electrode locations (markers); Cz, C1, CP6, CP4, FC6, FT8, and AF8. (D), (E) Average EF across all participants of the M1–IPL and M1–IFG groups, respectively. (F) Illusion-related activity (group effect) in the left M1, the right IPL and the right IFG during illusion with knowledge.

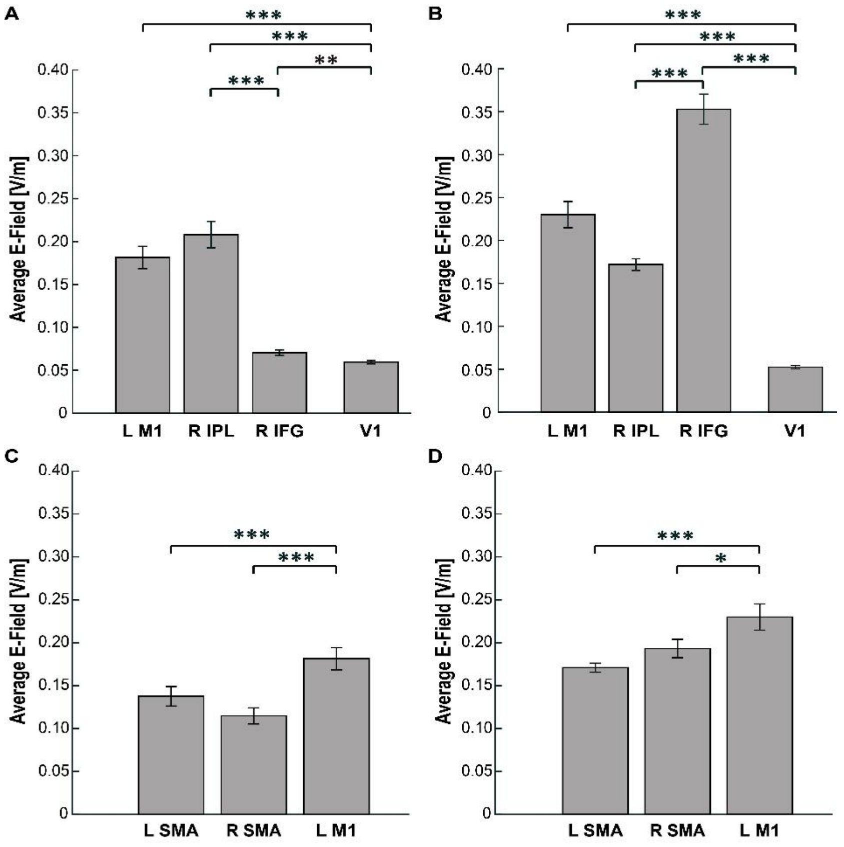


**Fig. S6.** Mean estimated EF strength in ROIs across participants. (A) The values of the left M1, right IPL, right IFG and V1 ROIs in M1–IPL group. (B) The values of the left M1, right IPL, right IFG and V1 ROIs in M1–IFG group. (C) The values of the left SMA, right SMA and left M1 in M1–IPL group. (D) The values of the left SMA, right SMA and left M1 in M1–IFG group. The values of the left M1 are identical to those shown in panels A and B. Error bars indicate standard errors of the mean across participants. Black asterisks indicate significant difference between ROIs after Bonferroni correction. * $p = 0.002$; ** $p = 0.0015$; *** $p < 0.001$. V1: primary visual cortex.

#### Spinocerebellar activity

The illusion-related activity across 63 participants of the M1–IPL, M1–IFG and sham groups were identified in the foot section of the left M1, the bilateral SMA, the right IPL, the right IFG, and the bilateral insular cortices (Figure S5F) as we observed in the fMRI experiment (Figure 2A). Within the predefined ROI (Table S2), the illusion-related activity (33 voxels) was identified in the right anterior cerebellar paravermis (peak coordinates x, y, z = 14, -38, -26), though the cluster size did not reach statistical significance. This region corresponds to the foot section of the spinocerebellum (4), which likely receives the kinesthetic (muscle spindle) signals from the right foot. We extracted individual brain activity (parameter estimates) from the cluster in each participant. Then, we calculated the mean and standard error of the mean (SEM) across participants in each group. A one-sample t-test revealed that the mean cerebellar activity across participants was significantly different from zero ($p = 0.017$ for M1-IPL group, $p = 0.007$ for M1-IFG group, $p = 0.001$ for sham group; Figure S6). No significant between-group differences were observed. Thus, the spinocerebellum likely received kinesthetic inputs during illusion with knowledge in the participants of the tDCS experiment, suggesting that the illusion with knowledge experienced by the participants of the tDCS experiment 1 is elicited not only through top-down processing but also through bottom-up processing.

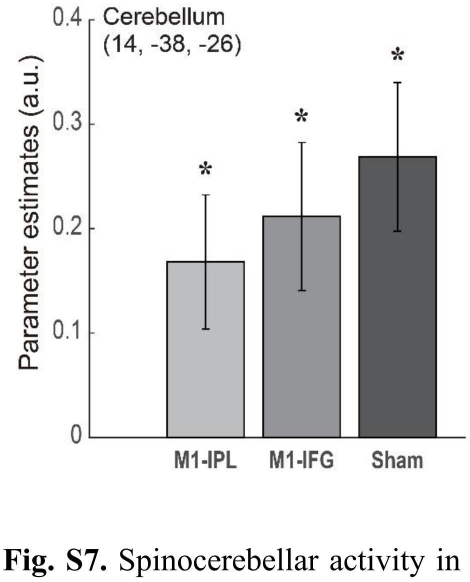


**Fig. S7.** Spinocerebellar activity in the participants of the tDCS experiment. Averaged brain activity (parameter estimate) across participants, extracted from the spinocerebellum cluster. Error bars indicate standard errors of the mean across participants. Black asterisks indicate significant difference from zero.

#### tDCS experiment 2

##### EMG recording and analysis

In the second tDCS experiment, we recorded electromyographic (EMG) activity using surface electrodes from the medial and lateral heads of the gastrocnemius, soleus and tibialis anterior muscle of the right leg during each trial. We measured EMG in all groups, however, since the results were similar across groups, we only reported EMG activity obtained from the M1–IPL group (Figure 7). After careful skin surface treatment, four pairs of electrodes (BA-U410 m(A)-015, Nihonsanteku Co., Ltd., Osaka, Japan) were placed over the belly of each muscle according to the Surface EMG for the Non-invasive Assessment of Muscles (SENIAM) Project (10). The ground electrode was placed over the right fibular head. The electrodes were attached to the cleaned skin using double-sided adhesive tape (Map-3M402, Nihonsanteku Co., Ltd., Osaka, Japan). Surface EMG (sEMG) signals were recorded using the EMG Multi Analysis Program MaP1038L (Nihonsanteku Co., Ltd., Osaka, Japan; sampling rate = 2048 Hz; high frequency filter = 1000 Hz; time constant = 0.03 sec). The EMG raw data were amplified (gain, 10000), and smoothed using a moving average. The root-mean-square (RMS) EMG values were calculated for each trial and each participant using the first 2–19 s of the 30–s resting period preceding the vibration (after this period, a countdown instruction to initiate the vibration was provided) and the first 2–13 s of the subsequent 15–s vibration period. The increase of EMG activity was determined by subtracting the value during the resting period from that during the tendon-vibration period. The mean of the EMG activity increase was calculated across participants for each trial (Figure 7). This was done for each muscle. We statistically evaluated if the EMG values are greater during the tendon-vibration period than during the resting period, however, no significant differences were found for any muscle in any trial.

After completion of the tendon-vibration trials, we also recorded EMG activity from the four muscles while the participant actually performed right ankle plantar flexion and maintained the maximal plantar flexion position for 15 seconds. The same analyses were conducted, and the results are shown in Figure 7. We confirmed significantly greater EMG values during the movement period than during the preceding resting period in all muscles. Although illusory plantar flexion can be experienced beyond the actual range of ankle motion (11), the EMG activity was negligible compared with that observed during actual movement.

**Inter-participant variability in illusion strength (tDCS effect)**

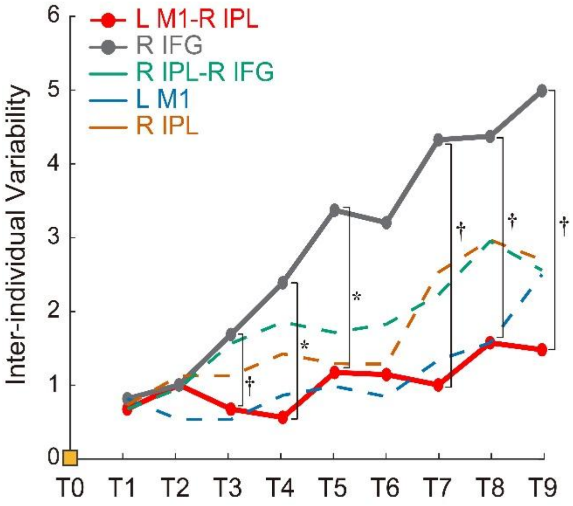


**Fig. S8**. Inter-participant variability in illusion strength (tDCS effect). We calculated inter-participant variability in illusion strength for each trial (from T1 to T9) in each tDCS group of the second tDCS experiment. Red solid line indicates the data obtained from the M1–IPL group, and grey solid line indicates the data from the IFG–alone group. Green dash line for IPL–IFG group, blue dash line for M1–alone group, and orange dash line for IPL–alone group. Levene's test revealed significant (and trend toward) difference in variance between the M1–IPL and IFG–alone groups (T3, $p = 0.08$; T4, $p = 0.01$; T5, $p = 0.01$; T7, $p = 0.07$; T8, $p = 0.05$; T9, $p = 0.09$). Black asterisks indicate significant difference in each trial between the M1–IPL and IFG–alone groups. IFG–alone group showed significantly greater inter-participant variability than M1–IPL group.
